# The fungal pathogen *Ustilago maydis* targets the maize corepressor TPL2 to modulate host transcription for tumorigenesis

**DOI:** 10.1101/2023.06.12.544564

**Authors:** Luyao Huang, Bilal Ökmen, Sara Christina Stolze, Melanie Kastl, Mamoona Khan, Daniel Hilbig, Hirofumi Nakagami, Armin Djamei, Gunther Doehlemann

## Abstract

*Ustilago maydis* is a biotrophic fungus that causes tumor formation on all aerial parts of maize. *U. maydis* secretes effector proteins during penetration and colonization to successfully overcome the plant immune response and reprogram host physiology to promote infection. In this study, we functionally characterized the *U. maydis* effector protein Topless (TPL) interacting protein 6 (Tip6). We found that Tip6 interacts with the N-terminus of ZmTPL2 through its two EAR (Ethylene-responsive element binding factor-associated amphiphilic repression) motifs. We show that the EAR motifs are essential for the virulence function of Tip6 and critical for altering the nuclear distribution pattern of ZmTPL2. We propose that Tip6 mimics the recruitment of ZmTPL2 by plant repressor proteins, thus disrupting host transcriptional regulation. We show that a large group of AP2/ERF B1 subfamily transcription factors are misregulated in the presence of Tip6. Our study suggests a regulatory mechanism where the *U. maydis* effector Tip6 utilizes repressive domains to recruit the corepressor ZmTPL2 to disrupt the transcriptional networks of the host plant.

## Introduction

*Ustilago maydis,* the causal agent of common corn smut, induces tumor formation in various aerial parts of maize (*Zea mays*). To establish successful infection, *U. maydis* employs a diverse array of effector proteins that facilitate interaction with the host plant and promote disease progression (Schuster et al., 2018). Previous studies have shown that *U. maydis* effectors exhibit organ-specific expression and adaptation during infection, as revealed by comparative transcriptome analysis of infected seedlings, adult leaves, and tassels (Skibbe et al., 2010). Among these effectors, nine have been identified as leaf-specific effectors, playing a crucial role in *U. maydis* virulence specifically in maize leaves (Schilling et al., 2014). For instance, See1 (Seedling efficient effector 1), the first characterized leaf-specific effector protein, interacts with the maize SGT1 protein, leading to the reactivation of host DNA synthesis and direct promotion of tumor formation in bundle sheath cells (Matei et al., 2018; Redkar et al., 2015). Additionally, Erc1 (enzyme required for cell-to-cell extension) has recently been found to have 1,3-glucanase activity, which is required for fungal cell-to-cell elongation in leaf bundle sheaths (Ökmen et al., 2022). Another leaf-specific effector whose deletion resulted in a significant reduction in tumors is UMAG_11060 (Schilling et al., 2014). However, its molecular function has yet to be clearly determined.

Regulating gene expression is crucial for plant responses to biotic and abiotic stress. The TOPLESS (TPL) and TOPLESS RELATED PROTEIN (TPR) co-repressors are involved in major plant hormone signaling, as well as meristem initiation and maintenance developmental pathways (Gallavotti et al., 2010; Liu et al., 2019; Long et al., 2006; Oh et al., 2014; Pauwels et al., 2010; Szemenyei et al., 2008). TPL/TPR proteins are recruited directly or indirectly by DNA-binding transcription factors to suppress the expression of target genes. Transcription factors typically interact with TPL through transcriptional repression motifs, such as the Ethylene-Responsive Factor (ERF)-associated amphiphilic repression (EAR) motif, characterized by the LxLxLx or DLNxxP sequence, which mediates interaction with TPL/TPR (Causier et al., 2012; Hiratsu et al., 2003; Kagale & Rozwadowski, 2011; Liu et al., 2019; Ohta et al., 2001). Notably, the APETALA2/Ethylene-Responsive Factor (AP2/ERF) superfamily of transcription factors was found to be enriched among the interactors of TPL/TPR proteins (Causier et al., 2012). This superfamily plays an important role in various aspects of plant growth and development, including response to biotic and abiotic stresses, hormone signaling, and pathogen defense (Chandler, 2018; Han et al., 2020; Koyama et al., 2013; Krishnaswamy et al., 2011; C. Wang et al., 2011). The AP2/ERF superfamily can be further classified into five subfamilies: Apetala 2 (AP2), Ethylene-Responsive Factors (ERF), Dehydration-Responsive Element-Binding Proteins (DREB), Related to Abscisic Acids-Intensive 3/viviparous 1 (RAV), and Soloist (Cheng et al., 2023; Sakuma et al., 2002).

ERF transcription factors normally interact with a cis-regulatory element known as the GCC box, which comprises the DRE/C-repeat (DRE/CRT) element and the ethylene-responsive element (ERE) (Büttner & Singh, 1997; Fujimoto et al., 2000; Hao et al., 1998; Masaru & Hideaki, 1995; Sessa et al., 1995). These factors are commonly found in the promoters of genes that respond to abiotic stress, jasmonate- and ethylene-inducible genes, and genes involved in pathogenesis (Büttner & Singh, 1997; Chakravarthy et al., 2003; Lorenzo et al., 2003; Maruyama et al., 2013; Masaru & Hideaki, 1995; Pré et al., 2008). The ERF family is categorized into six subgroups, designated as B1 to B6, with the B3 family being responsible for regulating multiple disease resistance pathway genes (McGrath et al., 2005; Moffat et al., 2012). Overexpression of AtERF1 resulted in the up-regulation of PLANT DEFENSIN1.2 (PDF1.2) and chitinases (Lorenzo et al., 2003; Solano et al., 1998). Similarly, the overexpression of the tomato ERF transcription factor Pti4 stimulated the expression of pathogenesis-related (PR) 1 and PR2, leading to increased resistance against fungal and bacterial pathogens (Gu et al., 2002). The B1 family plays a vital role in plant development, with the *Arabidopsis* gene *PUCHI* contributing to lateral root cell division and floral meristem identity (Bellande et al., 2022; Hirota et al., 2007; Karim et al., 2009). Homologs of PUCHI, such as BRANCHED SILKLESS1 (BD1) in maize and FRIZZY PANICLE (FZP)/BRANCHED FLORETLESS1 (BFL1) in rice, are also involved in floral meristem development (Chuck et al., 2002; Komatsu et al., 2003). Moreover, members of the B1 family, including AtERF3 and AtERF4, function as repressors that can downregulate the transcription level of reporter genes and the transactivation activity of transcription factors (Fujimoto et al., 2000; Ohta et al., 2000). TPL/TPRs serve as important targets for plant pathogens, and several *U. maydis* effectors that target maize TPL have been identified. Naked1 (Nkd1), a recently discovered effector, interacts with maize TPL through a C-terminal LxLxLx motif. This interaction results in the suppression of pathogen-associated molecular pattern (PAMP)-induced reactive oxygen species (ROS) bursts, leading to a significant decline in plant immunity (Navarrete et al., 2022). Moreover, the Nkd1-TPL interaction hinders TPL recruitment by the Aux/IAA repressor, resulting in the derepression of auxin signaling (Navarrete et al., 2022). The *U. maydis* cluster gene 6A encodes a family of five effector proteins known as Tips, which play a role in inducing auxin signaling (Bindics et al., 2022). Although neither of them contains an EAR domain, Tip1 and Tip4 compete with the Aux/IAA repressor for TPL binding, ultimately promoting the expression of auxin-responsive genes (Bindics et al., 2022). Additionally, the effector protein Jsi1, through its interaction with maize TPL, promotes the induction of jasmone/ethylene (JA/ET) (Darino et al., 2020). Unlike Nkd1, Jsi1 binds to the second WD40 domain of TPL/TPR proteins via a DLNxxP motif. This interaction enhances the biotrophic susceptibility of *Pst DC3000* in *A. thaliana* by upregulating genes associated with the ERF B3 branch of the JA/ET signaling pathway.

In this study, we report the functional characterization of the *U. maydis* effector gene *UMAG_11060*, which encodes the Topless interacting protein 6 (Tip6). Tip6 plays a crucial role in the full virulence of *U. maydis*, specifically inducing tumor formation upon seedling infection. We found that Tip6 interacts with maize ZmTPL2 through two EAR motifs, which are required for the virulence function of the effector. Furthermore, Tip6 binds to the N-terminus of ZmTPL2, which alters its subcellular localization. We propose that Tip6 disrupts the regulatory machinery of maize by recruiting TPL, thus interfering with the expression of maize transcription factors, particularly a group of AP2/ERF B1 subfamily proteins. These findings provide valuable insights into the mechanistic aspects of effector-mediated transcriptional regulation modulation in the host.

## Results

### Tip6 is a secreted effector protein

Tip6 is encoded by the *U. maydis* gene *UMAG_11060*, which was identified as an effector gene with an organ-specific virulence function in maize leaves (Schilling et al., 2014). To assess whether Tip6 is secreted during infection, we generated an *U. maydis* strain that secretes a Tip6-mCherry fusion protein under the control of its native promoter (SG200Δ*Tip6*_p*Tip6::Tip6*-mCherry) for subsequent confocal microscopy. To confirm secretion, we used a *U. maydis* strain carrying SG200Δ*pit2*_p*Pit2::Pit2*-*mCherry* as a positive control, which secretes a highly expressed apoplastic Pit2 effector (Mueller et al., 2013). A strain (SG200-p*Pit2*::*mCherry*) expressing cytoplasmic mCherry driven by the *pit2*-promoter served as a negative control for secretion. Confocal microscopy performed with infected maize leaves revealed that mCherry localizes inside the fungal hyphae, whereas Tip6-mCherry localizes at the periphery of the fungal hyphae, particularly at the hyphae tips, which is similar to the localization of Pit2-mCherry (Fig. 1A). To further examine secretion, we expanded the apoplastic space of infected maize cell by inducing plasmolysis through treatment with 1M NaCl. This showed that the secreted Tip6-mCherry and Pit2-mCherry, but not cytoplasmic mCherry, accumulate in the apoplastic space of infected maize cells (Fig. 1B).

**Fig. 1.**
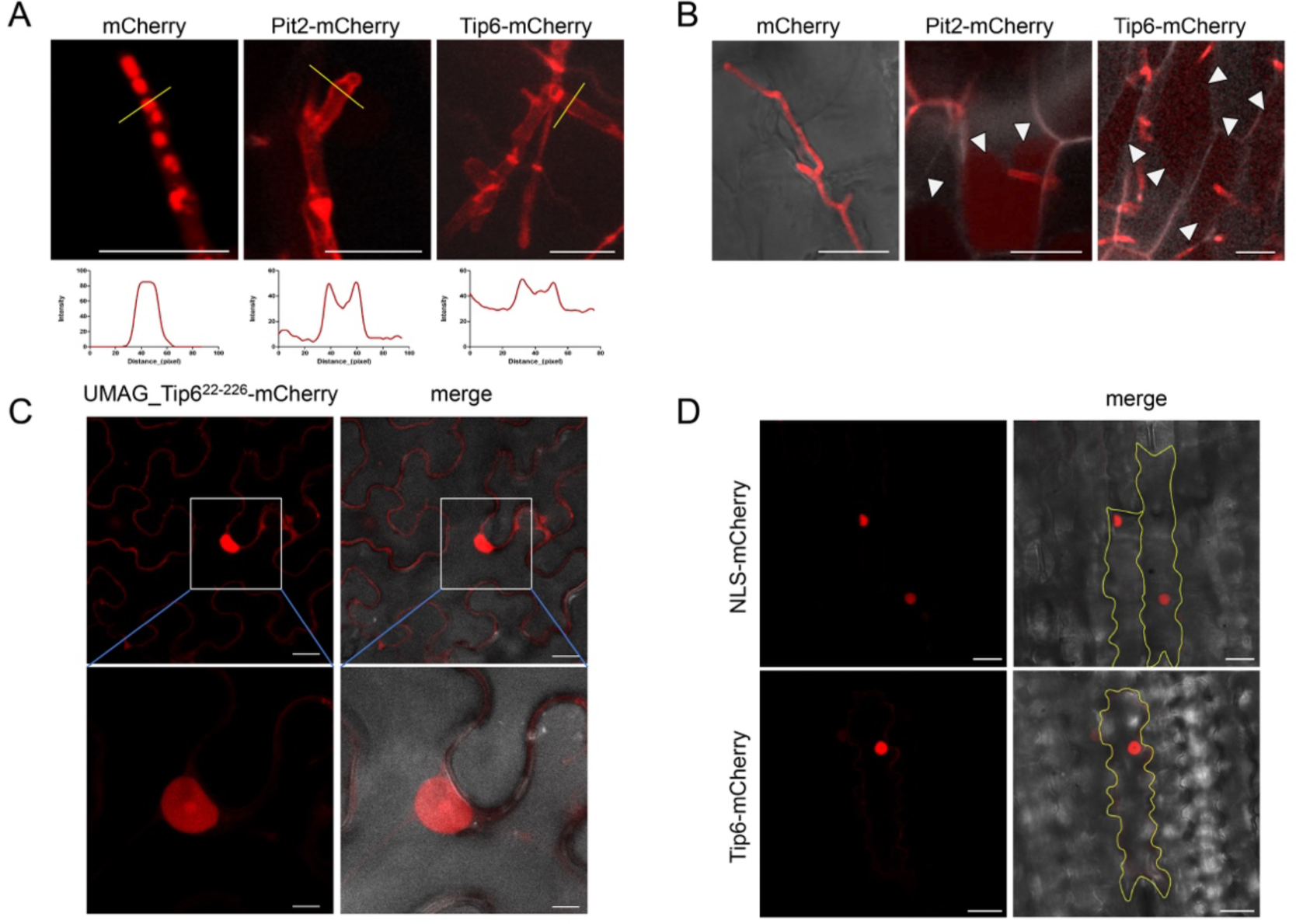
Tip6 is a secreted and translocated into the host nucleus and cytoplasm during the *U. maydis* infection. **A.** Confocal laser scanning microscopy images showing the localization of Tip6 in *U. maydis* infection at 3 dpi. Fluorescent signals of *U. maydis* strains expressing SG200-p*Pit2*::*mCherry*, SG200Δ*pit2*_p*Pit2*::*Pit2-mCherry*, or SG200Δ*Tip6*_p*Tip6*::*Tip6*-mCherry were visualized. SG200-p*Pit2*::*mCherry* localized inside the hyphae, while SG200Δ*pit2*_p*Pit2*::*Pit2-mCherry* and SG200Δ*Tip6*_pTip6::*Tip6-mCherry* mainly localized at the periphery of hyphae and hyphal tips. Fluorescence intensity profiles along the orange lines are displayed below the respective images. Scale bar, 20 μm. **B.** Secretion of Tip6 into the plant apoplast. Apoplastic spaces were enlarged by plasmolysis with 1 M NaCl. SG200-p*Pit2*::*mCherry* did not diffuse into the apoplast, while the mCherry signals of SG200Δ*pit2*_p*Pit2*::*Pit2-mCherry* and SG200Δ*Tip6*_pTip6::*Tip6-mCherry* diffused into the enlarged apoplast, as indicated by the arrows. Scale bar, 20 μm. **C**. Subcellular localization of Tip6 in *N. benthamiana* expressing p*2×35S::Tip6^22-226^-mCherry*. Images were observed at 2 dpi. Scale bar, 20 μm**. D**. Subcellular localization of Tip6 in maize epidermal cells expressing p*2×35S::NLS-mCherry* or p*2×35S::Tip6^22-226^-mCherry*. NLS-mCherry served as a nuclear marker. Images were taken 16-24 hours after transformation. Scale bar, 20 μm.

Next, we assessed Tip6 subcellular localization upon *Agrobacterium*-mediated expression in *Nicotiana benthamiana*. Tip6^22-226^-mCherry, lacking a predicted signal peptide, localized in both, the nucleus and cytoplasm of *N. benthamiana* (Fig. 1C). To verify the localization of Tip6 in maize, we examined the expression of Tip6^22-226^-mCherry in maize epidermal cells through biolistic bombardment. Confocal images showed that NLS-mCherry only localized in the maize nucleus, while Tip6^22-226^-mCherry localized in both, the maize nucleus and cytoplasm (Fig. 1D). In conclusion, our results show that Tip6 is secreted and translocated into the host nucleus and cytoplasm during *U. maydis* infection.

### Tip6 interacts with ZmTPL2

To elucidate the function of Tip6, we performed a co-immunoprecipitation (Co-IP) assay followed by mass spectrometry analysis to identify its potential host targets. First, we expressed Tip6^22–226^-GFP or GFP in *N. benthamiana* and conducted co-immunoprecipitation (Fig. S1A). Second, we infected maize seedlings with a Δ*Tip6* strain expressing a triple haemagglutinin (HA)-tagged Tip6 (SG200ΔTip6-p*Pit2::Tip6-3xHA*) or a triple HA-tagged mCherry with signal peptide driven by the *Pit2* promoter (SG200-p*Pit2::SP-mCherry-3xHA*), then performed immunoprecipitation (Fig. S1A&S1B). Results from the mass spectrometry analysis are listed in Tab. S1 and S2. Among the identified putative targets, the TOPLESS-RELATED 3 (NbTPR3) protein was one of the most significant in *N. benthamiana* samples. In maize samples, the maize TPL genes ZmTPL1, ZmTPL2, ZmTPL3 and ZmTPL4 were all identified as significant candidates, with ZmTPL2 being particularly noteworthy as it was only present in the Tip6 samples but absent in the controls (Fig. S1C).

To confirm the interaction between Tip6 and maize TPL family genes, we first conducted a yeast two-hybrid (Y2H) assay. The maize ZmTPL1, ZmTPL2, ZmTPL3, and ZmTPL4 were each fused to GAL4AD, while Tip6^22–226^ was genetically fused to GAL4BD. Our results revealed that Tip6 interacts with ZmTPL2 and ZmTPL3, as shown by the growth on high-stringency selection media (Fig. 2A). Conversely, no interaction was observed with ZmTPL1 and ZmTPL4 (Fig. 2A), indicating that Tip6 selectively interacts with ZmTPL2 and ZmTPL3. This suggests that Tip6 has a target specificity in maize TPL family genes.

**Fig. 2.**
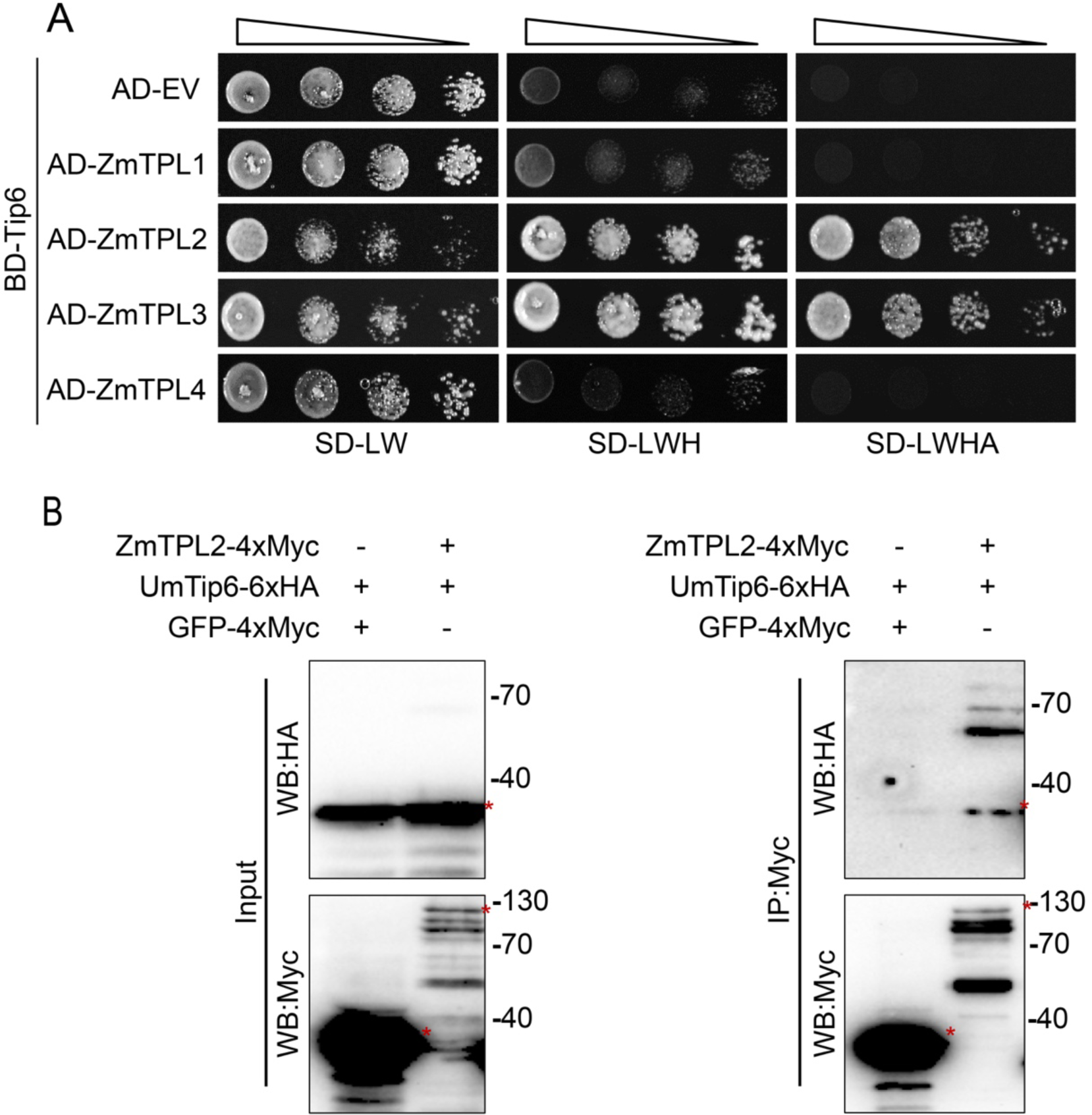
Tip6 interacts with ZmTPL2. **A.** The interaction between Tip6 and maize TPL was assessed by yeast two-hybrid assay. Yeast cells co-transformed with designated plasmids were dropped in ten-fold serial dilutions on synthetic defined (SD) medium plates lacking essential amino acids, including leucine and tryptophan (SD/-Leu/-Trp), histidine, leucine, and histidine (SD/-Leu/-Trp/-His) or leucine, tryptophan, histidine, and adenine (SD/-Leu/-Trp/-His/-Ade). Yeast cell growth on these plates was observed after 3-4 days. **B**. Confirmation of Tip6 and ZmTPL2 interaction by co-IP assay in *N. benthamiana*. *N. benthamiana* plants were transiently co-expressed with p*2×35S*::*Tip6^22-226^-6xHA* and either p*2×35S::ZmTPL2-4xMyc* or p*2×35S::GFP-4xMyc* (as a control) for 2 days. Proteins were extracted and immunoprecipitated using magnetic Myc-trap beads. The presence of input and IP proteins was determined using anti-HA or anti-Myc antibodies, respectively. Expected bands are indicated by red asterisks.

To further confirm the interaction between Tip6 and ZmTPL2, we performed a co-IP assay using *N. benthamiana*. The assay involved transient co-expression of Tip6^22–226^-6xHA with ZmTPL2-4xMyc, and as negative control GFP-4xMyc was co-expressed with Tip6^22–226^-6xHA. After *Agrobacterium* infiltration, proteins were extracted from *N. benthamiana* leaves and immunoprecipitated using α-Myc magnetic beads. The results revealed that Tip6^22–226^-6xHA co-immunoprecipitated with ZmTPL2-4xMyc, whereas GFP-4xMyc did not (Fig. 2B), confirming the specific interaction between Tip6 and ZmTPL2.

### Tip6 specifically interacts with the N-terminal domain of ZmTPL2

The TPL/TPR proteins have several highly conserved domains, with an N-terminal domain that consists of LisH, CTLH, and CRA domains, as well as a C-terminal domain consisting of two WD repeats (Ke et al., 2015; H. Ma et al., 2017; Martin Arevalillo et al., 2017). To elucidate, which domains of ZmTPL2 are involved in interaction with Tip6, we conducted Y2H assays with various truncated forms of ZmTPL2, such as the N-terminal domain (ZmTPL2^N^), C-terminal domain (ZmTPL2^C^), N-terminus lacking the CRA domain (ZmTPL2^NΔCRA^), and N-terminus CRA domain (ZmTPL2^CRA^) (Fig. 3A). The results showed a strong interaction between the N-terminus of ZmTPL2 and Tip6, indicating that ZmTPL2^N^ is responsible for binding to Tip6, but not the WD40 repeat domain (Fig. 3B).

**Fig. 3.**
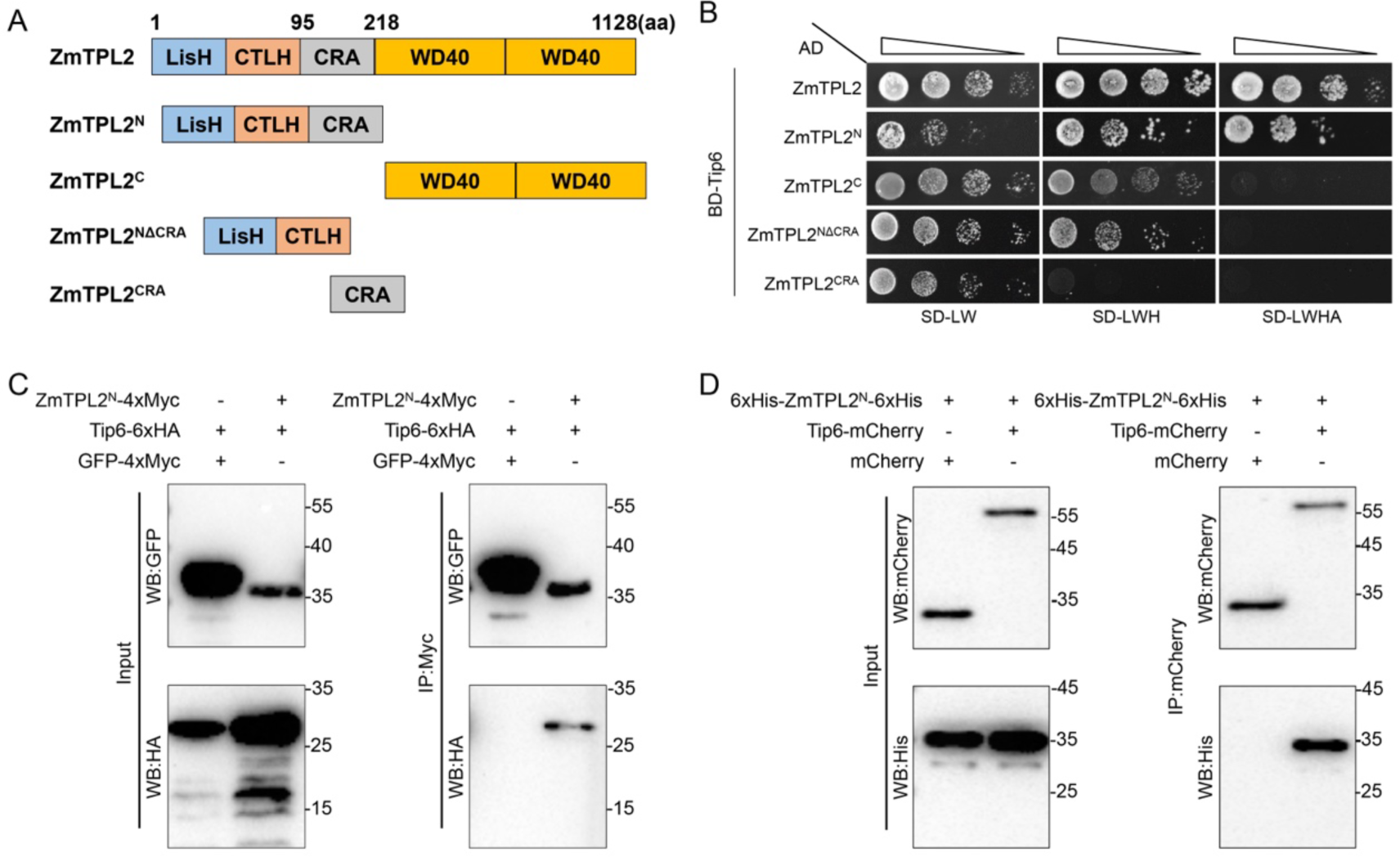
Tip6 interacts with the N-terminus of ZmTPL2. **A**. Schematic representation of ZmTPL2 and its truncated variants. The domain includes Lish (LIS1 homology domain), CTLH (C-terminal LisH motif domain), CRA (CT11-RanBPM domain), and the WD40 repeat domain. The variants include ZmTPL2^N^ (1-218 aa of ZmTPL2 N-terminal domain), ZmTPL2^C^ (216-1128 aa of ZmTPL2 C-terminal domain), ZmTPL2^NΔCRA^ (1-95 aa of ZmTPL2 N-terminus with Lish and CTLH domains), and ZmTPL2^CRA^ (90-218 aa of ZmTPL2 N-terminus, CRA domain). **B**. Interaction between Tip6 and ZmTPL2^N^ in the yeast two-hybrid assay. The plasmid carrying Tip6^22-226^ was co-transformed with plasmids carrying ZmTPL2 truncated variants into yeast cells. The Y2H assay was performed as described in Fig. 2A. **C**. Tip6 interaction with the N-terminal domain of ZmTPL2 was confirmed by a co-IP assay. *N. benthamiana* plants were transiently co-expressed with p*2×35S::Tip6^22-226^-6xHA* and either p*2×35S::ZmTPL2^N^-4xMyc* or p*2×35S::GFP-4xMyc* (control). The Co-IP assay was performed as described in Fig. 2B. Full-length bands are indicated by red asterisks. **D**.Tip6 interaction with the N-terminal domain of ZmTPL2 was demonstrated in an *in vitro* pull-down assay. Recombinant ZmTPL2^N^-His protein was mixed with mCherry or Tip6-mCherry, and protein immunoprecipitation was performed using magnetic mCherry-trap beads. The immunoprecipitated proteins were detected using anti-His and anti-mCherry antibodies.

To confirm the interaction between ZmTPL2^N^ and Tip6, we performed co-IP assays by co-expressing Tip6-4xMyc and ZmTPL2^N^-GFP or GFP-4xMyc in *N. benthamiana* leaves. This showed that Tip6 was precipitated with ZmTPL2^N^-GFP, but not with GFP-4xMyc **(Fig. 3C)**. Subsequently, we purified 6xHis-tagged ZmTPL2^N^ using Ni-NTA agarose and GST-Tip6^22-226^-mCherry and GST-mCherry using GST glutathione sepharose. We then removed the GST tag from Tip6^22-226^-mCherry or mCherry using prescission^®^ protease and separated each protein using size-exclusion chromatography (SEC). We collected and examined the proteins that migrated to the corresponding molecular masses **(Fig. S2)**. The purified His-ZmTPL2^N^-His and Tip6-mCherry or mCherry were subjected to an *in vitro* pull-down assay, which confirmed the interaction between ZmTPL2^N^ and Tip6 **(Fig. 3D)**. Together, these results strongly support a crucial role of ZmTPL2^N^ in the interaction with Tip6.

### Tip6 EAR motifs mediate its binding to ZmTPL2

The family of TPL/TPR proteins regulates gene expression in various biological processes through repression domains (RDs), such as the EAR domain (Causier et al., 2012; Hiratsu et al., 2003; Kagale & Rozwadowski, 2011; Liu et al., 2019; Masaru & Hideaki, 1995; Ohta et al., 2001). However, protein domain prediction using Pfam did not reveal any evidence of a repression domain in Tip6. We noticed a LxLxLx-type EAR motif sequence ‘LGLSLG’ in Tip6. To investigate the function of the ‘LGLSLG’ motif, we generated a Tip6_L92A mutant (the first leucine was replaced with alanine) and tested its interaction with ZmTPL2 using Y2H assays **(Fig. 4A)**. Our results showed that full-length Tip6 interacted with ZmTPL2, consistent with our previous findings. However, Tip6_L92A displayed a reduction in binding to ZmTPL2, suggesting that leucine 92 within the ‘LGLSLG’ motif is required for Tip6-ZmTPL2 interaction **(Fig. 4B)**. Therefore, we propose that Tip6 interacts with ZmTPL2 partially through this LxLxLx-type EAR motif.

**Fig. 4.**
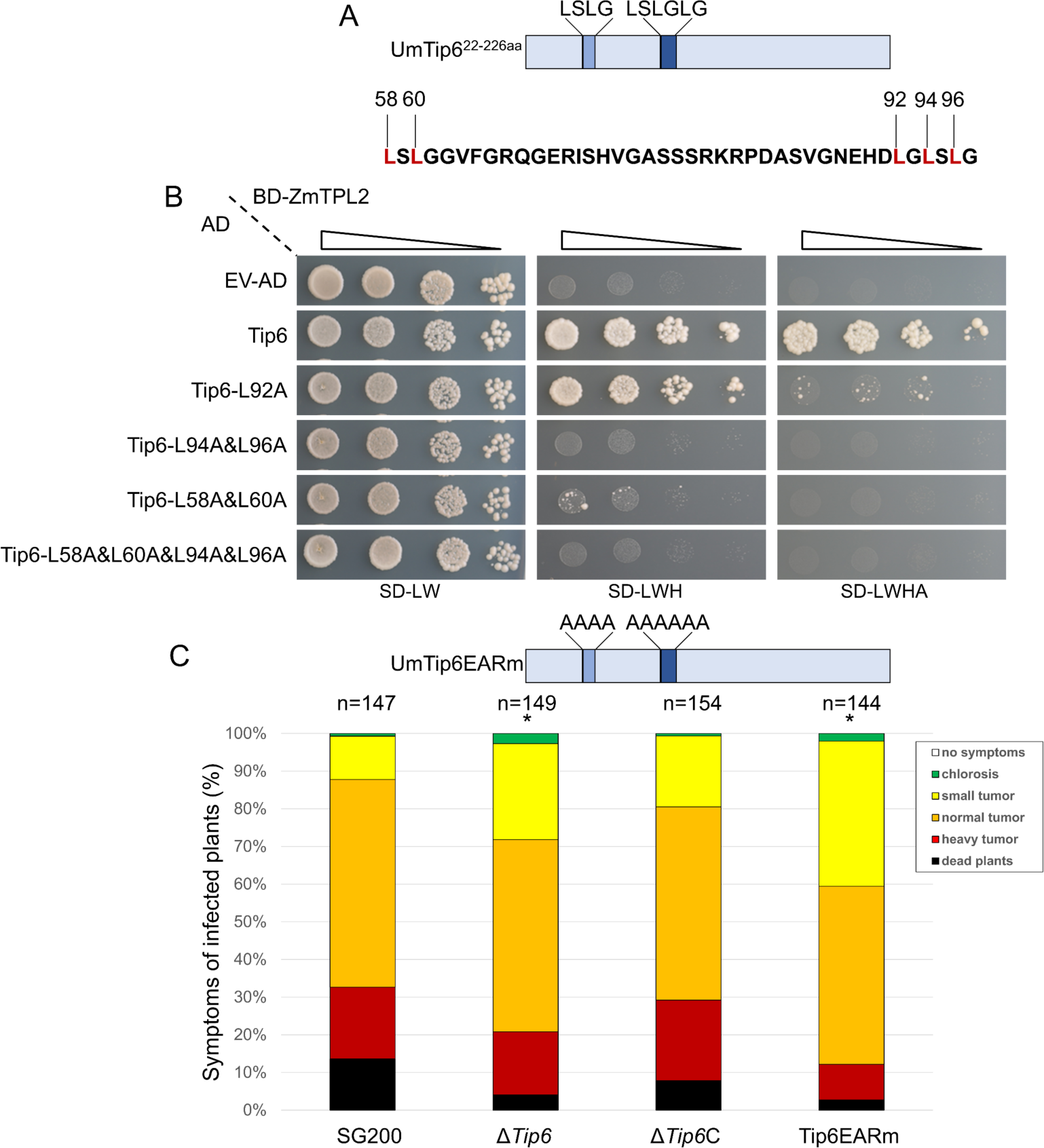
EAR motifs of Tip6 are essential for interaction with ZmTPL2 and virulence activity. **A.** Schematic representation of Tip6 EAR motifs. Tip6 contains two EAR repression domains, and the numbers above the protein sequence indicate the positions of the point mutations of the EAR motifs. **B**. Y2H assay to assess interactions between Tip6 EAR motif variants and ZmTPL2. The assay was conducted as described in Fig. 2A. **C**. Virulence activity of the Tip6 deletion mutant (Δ*Tip6*), its full-length complementation (Δ*Tip6*/C), and its EAR motifs mutant version (Tip6EARm). The protein sequence in the figure represents the mutated positions of the EAR motifs, which have been substituted with alanine. Maize seedlings (7 days old) were infected with *U. maydis* strains, and disease symptoms were scored at 12 dpi. Asterisks indicate significant differences (p < 0.05; Student’s T-test). The experiments were repeated three times.

After observing the remaining interaction between Tip6_L92A and ZmTPL2, we investigated the possibility of an additional repressive domain within Tip6. We identified a sequence ‘TELSLGG’ within Tip6, which we hypothesized to function as an EAR motif. To test this hypothesis, we generated the Tip6_L58A&L60A mutants, where leucine 58 and leucine 60 were substituted with alanine 58 and alanine 60. We conducted a Y2H assay with ZmTPL2 to determine the binding capability of Tip6_L58A&L60A **(Fig. 4A)**. Our results showed that Tip6_L58A&L60A exhibited insufficient growth on intermediate stringency medium and almost no growth on high stringency medium **(Fig. 4B)**. These findings confirm that the ‘TELSLGG’ sequence in Tip6 functions as an EAR motif, and it is crucial for Tip6 to bind to ZmTPL2.

To further confirm the importance of specific amino acids within the ‘LGLSLG’ motif, we generated the Tip6_L94A&L96A mutants and conducted a Y2H assay with ZmTPL2 **(Fig. 4A)**. The results demonstrated that Tip6_L94A &L96A failed to interact with ZmTPL2, as shown by almost no growth on the selection medium **(Fig. 4B)**. To assess the impact of both EAR motifs, we generated the Tip6_L58A&L60A&L94A&L96A mutants and tested their ability to bind ZmTPL2 **(Fig.4A)**. This showed that Tip6_L58A&L60A&L94A&L96A completely lost its interaction with ZmTPL2, as reflected by no growth on the plates (**Fig. 4B)**. These findings suggest that both EAR motifs within Tip6 are critical for its interaction with ZmTPL2.

In conclusion, our data provides evidence that Tip6 interacts with ZmTPL2 through two EAR motifs. The second and third leucine residues within the ‘LGLSLG’ EAR motif are particularly crucial for this interaction, while the first leucine is less important. Notably, this highlights the significance of the ‘TELSLGG’ motif in Tip6 for binding to ZmTPL2.

To assess the functional significance of the EAR motifs identified in Tip6, we conducted experiments to investigate their impact on the virulence of *U. maydis*. Specifically, we complemented a CRISPR/Cas9-generated Tip6 frameshift SG200 mutant with either the full-length Tip6 or an EAR mutant (Tip6EARm), in which the EAR motif sequence ‘LGLSLG’ and ‘LSLG’ were replaced with sequential alanine. Complementation with full-length Tip6 restored the virulence of Δ*Tip6* to a level similar to that of SG200. However, the Tip6EARm mutant could not restore the lost virulence and even exhibited a significant reduction in virulence compared to SG200 **(Fig. 4C)**. These findings strongly suggest that the identified EAR motifs, including the ‘LGLSLG’ motif and the ‘LSLG’ motif in Tip6 are crucial for the full virulence of *U. maydis*.

### Tip6 interferes with nuclear aggregation of ZmTPL2

To determine the subcellular localization of ZmTPL2, we transiently expressed ZmTPL2-GFP in *N. benthamiana* plants. Confocal imaging revealed that ZmTPL2-GFP accumulated in the nucleus and formed speckles **(Fig. 5 and Fig. S3A)**. Further investigation of ZmTPL2-GFP expression in maize epidermal cells showed a similar nuclear localization pattern to that observed in *N. benthamiana,* confirming the nuclear localization of ZmTPL2-GFP **(Fig. S3B)**.

**Fig. 5.**
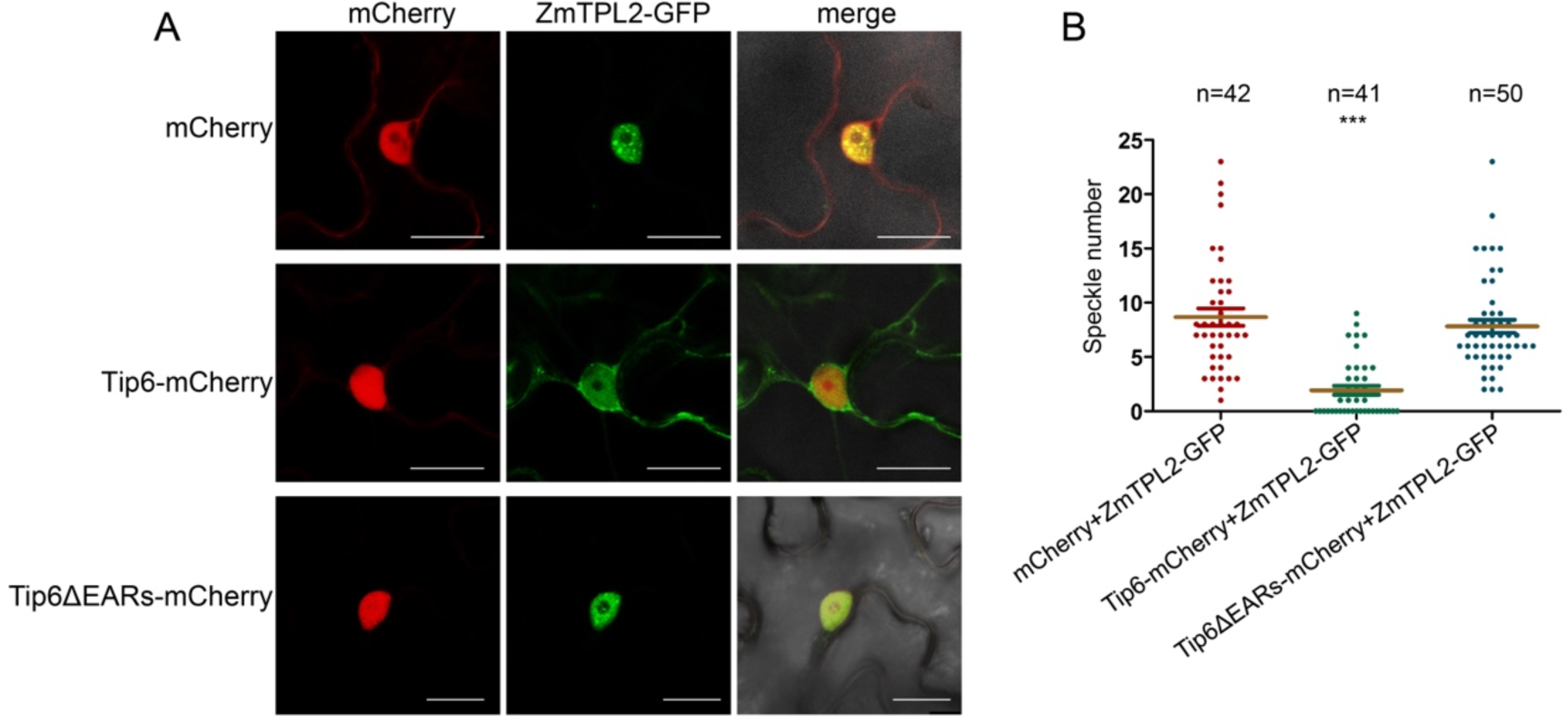
Effects of Tip6 on the nuclear distribution pattern of ZmTPL2 through its EAR motifs. **A.** Confocal microscopy images showing the nuclear distribution pattern of ZmTPL2 in the presence of mCherry, Tip6^22-226^-mCherry or Tip6^22-226^ΔEARs-mCherry. *N. benthamiana* epidermal cells were co-expressed with p*2×35S::ZmTPL2-GFP* and either p*2×35S::mCherry*, p*2×35S::Tip6^22-226^-mCherry*, or p*2×35S::Tip6^22-226^ΔEARs-mCherry* (with deleted EAR motifs). Scale bar, 20 μm. **B**. Comparison of the number of ZmTPL2 nuclear speckles in the presence of mCherry, Tip6^22-226^-mCherry or Tip6^22-226^ΔEARs-mCherry. The number of ZmTPL2 nuclear speckles was counted in the cells shown in Fig. 5A. Results were obtained from three independent experiments. N represents the total number of nuclei. (p<0.001, ANOVA, Tukeys).

To ascertain the effect of Tip6 on the subcellular localization of ZmTPL2, we co-expressed ZmTPL2-GFP with either Tip6-mCherry or mCherry in *N. benthamiana*. Remarkably, this showed that the presence of Tip6 resulted in a significant decrease in the number of nuclear speckles of ZmTPL2-GFP, whereas the speckle pattern remained unchanged in the presence of mCherry **(Fig. 5A)**. Additionally, it is worth noting that some fraction of ZmTPL2 is also localized in the cytoplasmic compartment **(Fig. 5A)**. As previously mentioned, the EAR motifs of Tip6 play a crucial role in binding to ZmTPL2 **(Fig. 4A)**. To examine the role of the EAR motifs in Tip6 in altering the localization of ZmTPL2, we co-expressed ZmTPL2-GFP with a Tip6ΔEARs-mCherry mutant lacking the EAR motifs in *N. benthamiana*. Surprisingly, confocal imaging revealed that the Tip6 mutant was unable to alter the nuclear distribution pattern of ZmTPL2, as seen by the ZmTPL2-GFP localized to nuclear speckles, similar to the case with mCherry co-expression **(Fig. 5A)**. Moreover, we found that Tip6ΔEARs cannot interact with ZmTPL2 via the Y2H assay **(Fig. S4)**. To further validate the role of Tip6 in altering the nuclear speckle formation of ZmTPL2, we quantified the number of speckles in *N. benthamiana* co-expressing cells. This showed that the number of ZmTPL2 speckles significantly decreased in cells co-expressing Tip6 compared to cells expressing mCherry. However, when we co-expressed Tip6ΔEARs mutant, the number of ZmTPL2 speckles was similar to that of mCherry **(Fig. 5B)**.

To investigate the impact of Tip6 on ZmTPL2 in maize, we transiently co-expressed ZmTPL2-GFP and either Tip6-mCherry or NLS-mCherry in maize epidermal cells. Confocal imaging revealed that both Tip6-mCherry and NLS-mCherry co-expression resulted in ZmTPL2-GFP localization with only a few speckles in the nucleus **(Fig. S5)**, which was unexpected as we hypothesized that Tip6 would decrease the number of speckles. In summary, these results demonstrate that Tip6 has an impact on the nuclear distribution pattern of ZmTPL2, and the EAR motifs of Tip6 are crucial in this process.

### Tip6 interferes with ZmTPL2-regulated transcription factors

To gain insights into the mechanisms underlying Tip6 virulence and assess the impact of its EAR motifs on infection, we performed transcriptome RNA-sequencing (RNA-seq) analysis on maize seedling leaves infected with SG200, Δ*Tip6*, or Tip6EARm at 3 days post-infection (dpi). Using a threshold of absolute log2Foldchange > 1 and *p*-value < 0.05, we found that compared to maize infected with SG200, Δ*Tip6* had 91 up-regulated and 71 down-regulated differentially expressed genes (DEGs), while Tip6EARm displayed 191 up-regulated and 115 down-regulated differentially expressed genes **(Fig. S6A and S6B, Tab. S3)**.

To examine the biological pathways associated with the DEGs, we performed a gene ontology (GO) enrichment analysis. The GO analysis of DEGs between Δ*Tip6* and SG200 revealed the regulation genes related to ‘cellular biosynthetic’, ‘transcription’, ‘gene expression’, and ‘RNA metabolic’ processes **(Fig. S6C, Tab. S3)**. Particularly, the majority of the regulated genes were involved in transcriptional regulation. Among the 18 transcription factors DEGs, nine belonged to the AP2/ERF family, with seven up-regulated genes belonging to the ERF B1 family. The remaining two down-regulated genes were classified as members of the AP2 and DREB families, respectively **(Fig. 6A)**. Among these DEGs, we observed the down-regulation of *Branched silkless 1* (*bd1*), a gene known to regulate inflorescence formation from spikelet formation in maize (Chuck et al., 2002). Additionally, we found that the maize genes *dbp4* (ZmDBP4) and *CBF3* were up-regulated, which were highly activated by cold and play regulatory roles in the abiotic stress responses of plants (Han et al., 2020; C. Wang et al., 2011). The down-regulated *ereb26*, a member of the AP2 subfamily, is known to be highly involved in floral formation (Kunst et al., 1989).

**Fig. 6.**
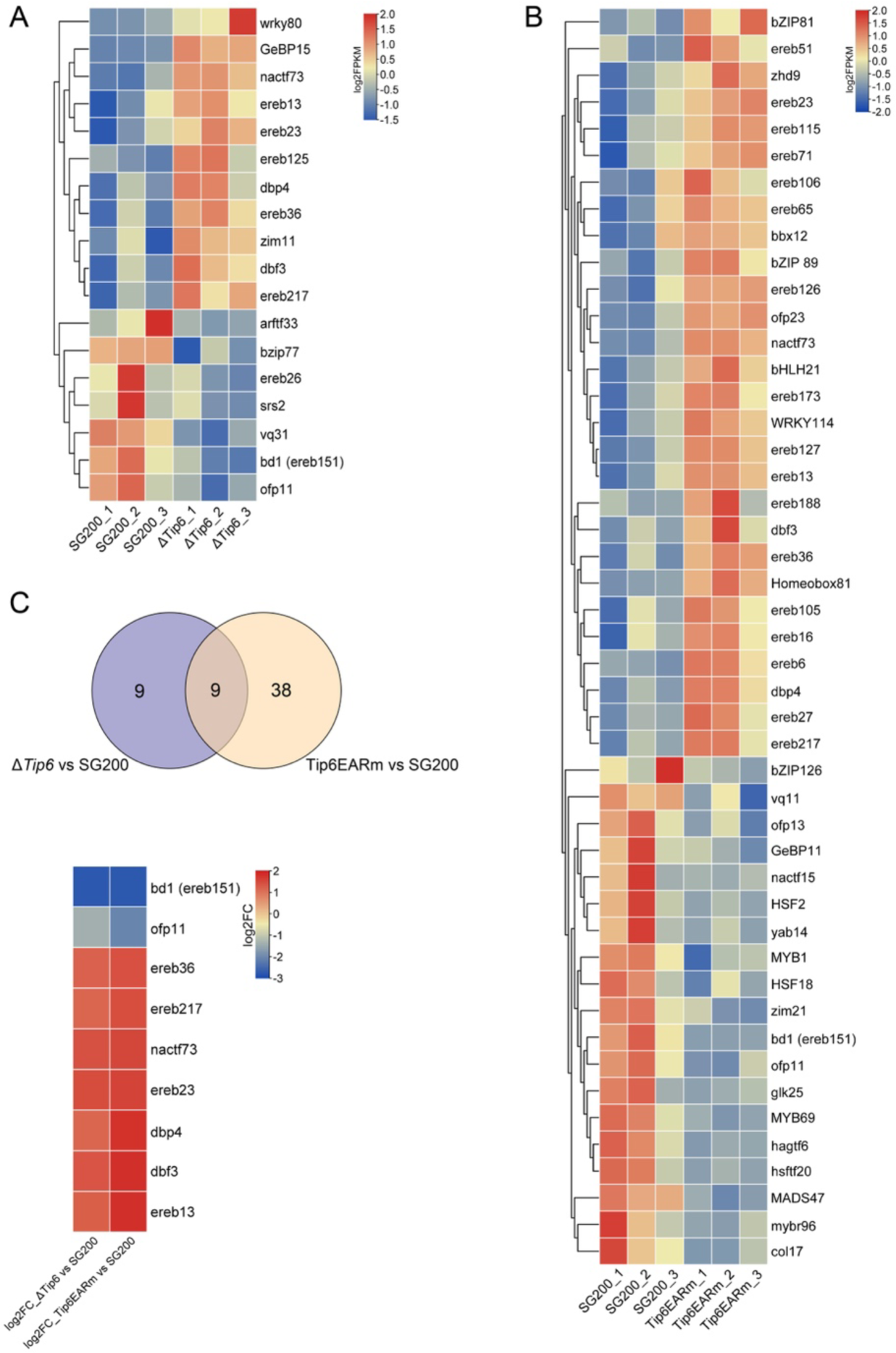
Impact of Tip6 on host transcription factors’ expression. **A.** Heatmap illustrating the DEGs in maize infected with Δ*Tip6* and SG200 strains. The log2FPKM values, based on the original FPKM values, were used to construct the heatmap. Rows and columns were created using the Euclidean distance method and the complete linkage method via TBtools (Chen et al., 2020). Gene expression is displayed on a red-to-blue spectrum, with red indicating high expression and blue indicating low expression. **B**. Heatmap of DEGs in maize infected with Tip6EARm and SG200 strains. The heatmap was generated similarly to Fig .6A. **C**. Venn diagram depicting the shared DEGs in Δ*Tip6* and Tip6EARm compared to SG200 samples. **D**. Heatmap showing the shared DEGs in Δ*Tip6* versus SG200 and Tip6EARm versus SG200. The log2fold change in expression of Δ*Tip6* and Tip6EARm compared to SG200 is depicted.

Several of the identified genes in the comparison of Δ*Tip6* and SG200 have known orthologs in other species. For instance, ovate transcription factors have been shown to be repressor regulators in *Arabidopsis* ovate family protein 1 (AtOFP1), which inhibits cell elongation, and rice ovate family protein 6 (OsOFP6), which regulates lateral root growth, leaf inclination and responses to abiotic stimuli (Y. Ma et al., 2017; Sun et al., 2020; S. Wang et al., 2007). SHI/STY transcription factors are involved in organ development and hormone regulation (Fang et al., 2023; He et al., 2019; Zhang et al., 2015). The *Arabidopsis* homologue of ZmNAC73, Jungbrunnen1 (JUB1), acts as a negative regulator of leaf senescence (Wu et al., 2012). Additionally, we observed the up-regulation of *wrky80*, which belongs to a family of transcription factors known to play crucial roles in disease resistance and response to abiotic stress (Hu et al., 2021; Rushton et al., 2010). Overall, Tip6-responsive maize genes are involved in various biological processes, including meristem maintenance, floral organ morphogenesis, disease resistance, and abiotic stress response.

The GO analysis of DEGs between Tip6EARm and SG200 showed the regulation of genes associated with ‘cellular biosynthetic’, ‘transcription’, ‘gene expression’, and ‘RNA metabolic’ processes **(Fig. S6D, Tab. S3)**. Among the 48 differentially expressed transcription factors, 20 belonged to the AP2/ERF family. Interestingly, 19 of these AP2/ERF family members were up-regulated, while *Branched silkless 1* (*bd1*) showed down-regulation, as seen in Δ*Tip6*-infected maize **(Fig. 6B)**. Additionally, the comparison of DEGs between Δ*Tip6* and SG200, as well as between Tip6EARm and SG200, revealed that 9 DEGs exhibited similar regulation, with 7 of them belonging to the AP2/ERF family **(Fig. 6C and D)**. These findings suggest that Tip6 may suppress a specific group of host transcription factors, particularly those in the AP2/ERF family, suggesting its regulatory role in the transcriptional modulation of the host plant.

We therefore hypothesized that the interaction between Tip6 and ZmTPL2 affects the recruitment of host transcription factors and subsequent host gene expression. However, we did not find any differentially expressed maize TPL genes in our RNA-seq data. To identify potential regulators of the DEGs, we performed an enrichment analysis using the PlantTFDB database (http://planttfdb.cbi.pku.edu.cn) (Jin et al., 2017; Tian et al., 2020). In the comparison of Δ*Tip6* vs. SG200 DEGs, we observed enrichment of 11 transcription factors (p<0.05). Notably, *ereb125* exhibited up-regulation and putative regulation of 31 DEGs **(Fig. S7A)**. Furthermore, *ereb125* was implicated in the putative regulation of other up-regulated transcription factors, including *dbp4*, *zim11*, *NAC73*, and *wrky80*.

Similarly, in the comparison of Tip6EARm vs. SG200 DEGs, we observed enrichment of 31 transcription factors, including ZmTPL3 interactors *ramosa2* and *ereb147* (Liu et al., 2019) Moreover, *ereb125* was found to putatively regulate 61 DEGs, including *dbp4*, *NAC73*, *ereb51*, *hb81*, *bHLH21*, and *ereb127* **(Fig. S7B, Tab. S3)**. These findings strongly suggest that Tip6 plays a significant role in the regulation of host transcription factors, particularly those belonging to the AP2/ERF family.

## Discussion

In this study, we demonstrate that the *U. maydis* effector Tip6 promotes tumorigenesis by targeting maize ZmTPL2 through its EAR motifs. Tip6 mimics the plant recruitment mechanism, disrupting host gene expression by recruiting the N-terminal domain of ZmTPL2 via its EAR domains. Mutation of the leucine residue in the EAR motifs abolishes the interaction of Tip6 with ZmTPL2. Moreover, Tip6EARm fails to restore the defective virulence phenotype of Δ*Tip6*. Interestingly, the *U. maydis* effector Nkd1 also interacts with ZmTPL2 via an EAR motif, but deletion or mutation of the EAR motif in Nkd1 reduces the interaction without affecting its ability to complement Δnkd1-deficient virulence, unlike Tip6 (Navarrete et al., 2022).

Tip6 binding to ZmTPL2 is diminished upon mutation or deletion of either of the EAR motifs (sequence with LSLG or LGLSLG), indicating the importance of having two EAR motifs for enhanced binding capacity. This finding raises the question of why Tip6 possesses two EAR motifs. To shed light on this, we can draw a parallel with the Aux/IAA repressor 7 (IAA7) in *Arabidopsis*, which also has two EAR motifs with different roles. In AtIAA7, the second EAR motif plays a minor role in repression but is crucial for interaction with TPR1, whereas the first EAR motif plays a major role in both repression and interaction with all TPL/TPR members (Lee et al., 2016). This suggests that the presence of two EAR motifs in Aux/IAA repressors provides greater repression capacity and interaction diversity compared to those with a single EAR motif (Lee et al., 2016). Similarly, the recently discovered rhizogenic *Agrobacterium* protein RolB has an N- and a C-termini EAR motif, but only the C-terminal EAR motif is required for TPL recruitment and hairy root development (Gryffroy et al., 2023). By considering this, we can speculate that the presence of two EAR motifs in Tip6 may confer similar functional advantages. Although the exact mechanism and significance of this enhanced binding capacity in Tip6 remain unclear, our findings demonstrate that Tip6EARm, similar to the Nkd1SRDX mutants with increased binding capacity to TPL, results in a significant loss of virulence. These findings highlight the critical role of the precise binding strengths exhibited by Tip6 or Nkd1 to ZmTPL2 in their pathogenic role in *U. maydis*.

Our RNA-seq analysis identified 57 differentially expressed maize transcription factors in the presence of Tip6. Among the 22 AP2/ERF DEGs, 20 were downregulated, with 13 belonging to the B1 family and 2 being upregulated. Jsi1 and RolB activate the ERF B3 branch of the JA/ET signaling pathway by recruiting TPL (Darino et al., 2020; Gryffroy et al., 2023). The ERF B3 subfamily acts as a positive transcriptional regulator, activating defense-related genes in the JA/ET hormone signaling pathways. Interestingly, we did not find any B3 group genes among our data, and GO analysis of Tip6-regulated DEGs shows that they are not enriched in the JA/ET pathway, which is in line with different ERF branches being affected. Three pairs of paralogues were found in the B1 branch genes of DEGs: dbf3/ereb36, ereb105/ereb16, and ereb13/ereb217 (Cheng et al., 2023). Therefore, we propose that the ERF branch affected by Tip6 is more closely related to plant growth and development, influencing leaf tumor formation by *U. maydis*.

Our study highlights *ereb125* as a potential regulator of a significant number of DEGs. Although the exact function of *ereb125* in maize is yet to be determined, its ortholog in *Arabidopsis*, *LEAFY PETIOLE* (*LPE*) (AT5G13910.1) was initially discovered in mutant screens for leaf development (van der Graaff et al., 2000). In wild-type *Arabidopsis*, the expression of the *LPE* gene is very low and strongest in young leaves, with weaker expression in older leaves, suggesting tissue-specific roles (van der Graaff et al., 2000). These findings suggest a potential role for *ereb125* in leaf development. Therefore, we hypothesize that the interaction between Tip6 and ZmTPL2 may involve *ereb125*, thus promoting leaf tumor formation.

*U. maydis* effector genes display highly distinct stage-specific expression patterns during maize infection, suggesting differences in their functions and host adaptation (Lanver et al., 2018). Among these effectors, Tip6 shows peak expression at 2 dpi during early biotrophic development, while Jsi1 peaks at 4 dpi, and Nkd1 at 6-8 dpi during tumor formation (Darino et al., 2020; Navarrete et al., 2022). Tip1-Tip5, except for Tip3, which peaks at 4 dpi, all peak at 2 dpi (Bindics et al., 2022). This time-staggered expression pattern suggests not only a potential strategy to prevent competitive TPL recruitment, but also functional differences among these effectors. These effectors target TPL, exerting pleiotropic effects. Jsi1, Nkd1, and Tip6 recruit TPL through repression domains (LxLxLx or DLNxxP), while Tip1-Tip5 recruit TPL in a repression domain-independent manner (Bindics et al., 2022). Furthermore, Jsi1, Nkd1, and Tip1-Tip5 activate genes involved in hormone signaling pathways, including JA/ET, auxin, and SA, by relieving TPL-mediated repression and promoting plant defense responses, thereby ultimately increasing susceptibility to *U. maydis* infection (Bindics et al., 2022; Darino et al., 2020; Navarrete et al., 2022). However, Tip6 appears to have different regulatory effects, potentially influencing plant development during leaf tumor formation by interfering with ZmTPL2. These findings suggest the diverse strategies employed by *U. maydis* effectors to target and impact TPL activity, emphasizing the critical role of TPL as a central hub of plant transcriptional regulation during *U. maydis* invasion. Similar mechanisms have been observed in other species, such as the oomycete *Hyaloperonospora arabidopsidis* (Hpa) effector HaRxL21 and the rhizogenic *Agrobacterium* protein RolB, both of which interact with TPL through their EAR motifs to regulate host genes (Gryffroy et al., 2023; Harvey et al., 2020). Our study demonstrates that Tip6 recruits the N-terminus of ZmTPL2 through its two EAR motifs. This changes the formation of nuclear speckles and interferes with the dysregulation of many host transcription factors. Notably, we observed a down-regulation of ERF B1 branch genes associated with host development. In conclusion, we propose a simplified working model illustrating the interference of Tip6 with ZmTPL2 **(Fig. 7)**. We hypothesize that this mechanism may directly contribute to the specific virulence of Tip6 in leaf tumor formation. Our study sheds light on the function of Tip6 through protein interactome and gene transcriptome analysis. Future studies will aim to fully comprehend the molecular mechanisms by which Tip6 manipulates ZmTPL2 and influences leaf tumor formation.

**Fig. 7.**
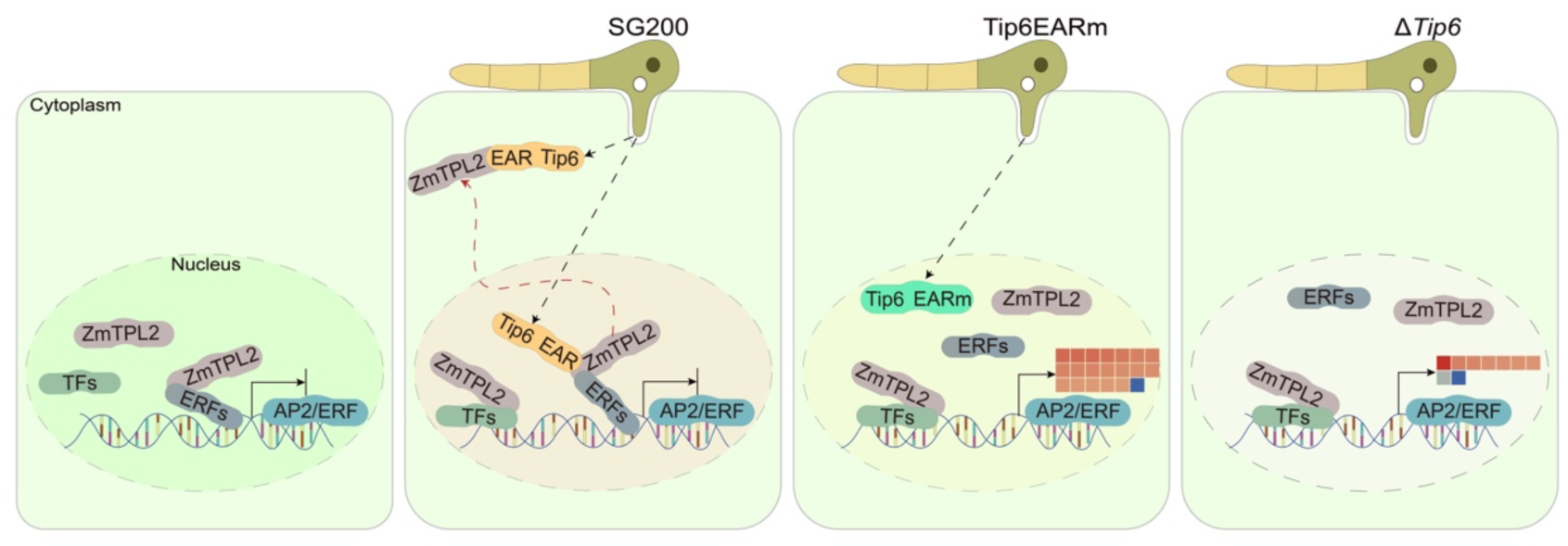
Tip6 impacts host gene regulation by interacting with the ZmTPL2 corepressor. The healthy plant cell where ERF family transcription factors recruit ZmTPL2 and repress AP2/ERF family transcription factors. While some unknown transcription factors (TFs) present, they do not recruit ZmTPL2 yet. SG200 strain infected cell, where Tip6 is expressed and secreted into the plant host. Tip6 interacts with ZmTPL2 utilizing EAR repressive motifs, leading to a change in the localization of ZmTPL2 from the nucleus to the cytoplasm and interfering with its repressor proteins. Tip6 with the EAR motif mutated into Alanine, which loses the ability to interact with ZmTPL2.This loss of interaction causes an unknown impact that leads to the mis-regulation of AP2/ERF family transcription factors. The knockout mutant, which lacks Tip6 and its interaction with ZmTPL2, suggesting that Tip6 plays a crucial role in the pathogenesis of *U. maydis*.

## Material and Methods

### Strains and plants growth conditions

Virulence assays were performed using the solo-pathogenic *U. maydis* strain SG200 and its derivative strains. *U. maydis* strains were grown in YEPS light liquid medium (0.4% yeast extract, 0.4% peptone and 2% sucrose) at 28°C with agitation at 200 rpm or on Potato Dextrose Agar (PD; Difco) plates at the same temperature. Plasmid vector cloning utilized *Escherichia coli* Top10 strains, while heterologous *in vitro* protein expression was performed using the *E. coli* BL21 (DE3) strain. Transient heterologous protein expression and protein subcellular observation in *N. benthamiana* were performed using the *Agrobacterium tumefaciens* GV3101 strain. All bacterial strains were cultured in dYT liquid medium or YT agar plates supplemented with suitable antibiotics. The yeast two-hybrid (Y2H) assay employed the AH109 yeast strain, which was cultivated in YPD liquid medium or SD agar plates lacking appropriate amino acids at 30°C.

*Zea mays* Golden Bantam (GB) plants were grown in a greenhouse under controlled conditions of 16 hours of light at 28°C and 8 hours of darkness at 22°C. *N. benthamiana* plants were cultivated in a growth chamber with 16 hours of light and 8 hours of darkness at 22°C. Information of the primers used in this study is provided in **Tab. S4**.

### Subcellular localization in tobacco and maize

The coding sequence of Tip6 lacking the signal peptide was fused with mCherry fluorescence tags, while the coding sequence of ZmTPL2 was fused with a GFP fluorescence tag. These constructs were transiently expressed in *N. benthamiana* leaves through *Agrobacterium*-mediated transformation. Fluorescence signals were visualized and captured using a Leica SP8 confocal laser scanning microscope (Leica, Germany). For maize, six-day-old leaves were bombarded with gold particles coated with the respective plasmids, as previously described in the protocol (Djamei et al., 2011). Confocal microscopy was performed on the samples after a 16–24-hour transformation.

### Yeast two-hybrid assay

The yeast two-hybrid assay was performed according to the Clontech Yeast Protocol Handbook. The coding sequence of Tip6, along with its variants were cloned into the pGBDT7 vector as bait. The encoding sequences of ZmTPL2 and its truncated variants were cloned into either the pGBDT7 or pGADT7 vectors. The resulting pGADT7 and pGBDT7 constructs were transformed into the yeast strain AH109, and the transformed cells were screened on nutrient-deficient plates under different stringency conditions: low stringency (SD-Leu-Trp), medium stringency (SD-Leu-Trp-His), and high stringency (SD-Leu-Trp-Ade-His). The plates were then incubated at 28 °C for 4-5 days.

### Co-immunoprecipitation and mass spectrometry analysis

Seven-day-old maize seedling leaves were infected with *U. maydis* carrying either SG200*ΔTip6-Ppit2::Tip6-3xHA* or SG200-*Ppit2::SP-mCherry-3xHA*. The infection assays were performed as previously described (Redkar & Dohlemann, 2016). At 3 dpi, the infected leaves (four biological replicates per sample) were collected and ground into powder using liquid nitrogen. For protein extraction, 500 uL of the ground powder was mixed with 1.5 mL of lysis buffer (50 mM Tris-HCl pH 7.5, 150 mM NaCl, 2 mM EDTA, 10% glycerol, 1 % [v/v] Triton X-100, 5 mM dithiothreitol (DTT) and protease inhibitor cocktail (Rothe. Cat. No. 04693159001)). After incubating on ice for 30 min with inversions every 10 min, the protein supernatant was obtained by centrifugation twice at 13,000 rpm 4 °C for 30 min.

Similarly, six-week-old *Nicotiana benthamiana* plant leaves were infiltrated with *Agrobacterium* carrying either *p2×35S::GFP* or *p2×35S::*Tip^22-226^-GFP. At 3 dpi, the infiltrated leaves (three biological replicates per sample) were harvested and ground into powder using liquid nitrogen. The ground powder was mixed with the same protein extraction buffer used in the maize samples and incubated on ice for 30 min with inversion every 10 min. The samples were centrifuged twice at 4°C for 30 min at 13,000 rpm, and the resulting supernatant was transferred to a new tube.

For both samples, 1 mL of the protein supernatant was mixed with 10 μL of respective antibody-coated beads (anti-HA magnetic beads (Thermo Fisher, Cat. No. 88836) for maize samples, and anti-GFP beads (ChromoTek GFP-Trap Agarose Cat. No. gta) for *N. benthamiana* samples). After incubating at 4°C for 1 h with constant rotation, the protein-bound beads were washed three times with extraction buffer and three times with extraction buffer without Triton X-100 before being subjected to mass spectrometry analysis.

Next, for the maize samples, the beads were re-dissolved in 25 µL digestion buffer 1 (50 mM Tris, pH 7.5, 2 M urea, 1 mM DTT, 5 ng/µL trypsin) and incubated at 30 °C in a thermomixer at 400 rpm for 30 min. The beads were then pelleted, and the supernatant was transferred to a fresh tube. To the beads, 50 µL of digestion buffer 2 (50 mM Tris, pH 7.5, 2 M urea, 5 mM CAA) was added. After mixing the beads, they were pelleted again, and the supernatant was collected and combined with the previously obtained supernatant. The combined supernatants were incubated overnight at 32 °C in a thermomixer at 400 rpm, while being protected from light. The digestion was stopped by adding 1 µL of TFA, and desalting was performed using C18 Empore disk membranes following the StageTip protocol (Rappsilber et al., 2003).

For the *N. benthamiana* samples, the beads were eluted with SDT buffer (4% SDS, 100 mM Tris-HCl pH 7.6, 0.1 M DTT). The SDS-eluted proteins were then subjected to a filter-aided sample preparation protocol (FASP) (Wiśniewski et al., 2009). Initially, the samples were reduced by adding 1/10 volume of 1 M DTT (100 mM final) and incubated at 95 °C for 5 min. Next, the samples were diluted with 8 M urea in 0.1 M Tris/HCl pH 8.5 (UA) to achieve a compatible detergent concentration (maximum 0.5% SDS). The diluted samples were loaded onto filters (Sartorius, Vivacon 500, VN01H22, 30 kDa cutoff) by centrifuging at 14,000 g for 10 min. Following that, the samples were washed with UA by centrifugation at 14,000 g for 10 min and alkylated using 100 µL of 55 mM chloroacetamide in UA. The mixture was then incubated in the dark for 20 min, followed by centrifugation at 14,000 g for 10 min. After three washes with UA, the filter was transferred to a new Eppendorf tube. Next, 50 µL of LysC solution (stock: 0.5 µg/µL Lys-C (WAKO) in 50 mM NH_4_HCO_3_ (ABC), working solution: dilute with UA to an enzyme: protein ratio of 1:100) was added, and the sample was mixed and incubated for 3 h at room temperature (RT). Subsequently, 300 µL of trypsin solution (stock: 1 µg/µL in 1 mM HCl, working solution: dilute with UA to an enzyme: protein ratio of 1:100) was added, and the samples were mixed and incubated overnight at RT. After centrifugation, 50 µL of ABC were added, and the samples were centrifuged at 14,000 g for 10 min. The resulting flow-through containing the peptides was acidified with TFA to a final concentration of 0.5%. Desalting of the flow-through was performed using stage tips with C18 Empore disk membranes (3 M) following the protocol described previously (Rappsilber et al., 2003).

For LC-MS/MS data acquisition, dried peptides were re-dissolved in a solution containing 2% ACN, 0.1% TFA for analysis and adjusted to a final concentration of 0.1 µg/µl. Samples were analyzed using an EASY-nLC 1200 (Thermo Fisher) coupled to a Q Exactive Plus mass spectrometer (Thermo Fisher). Peptides were separated on 16 cm frit-less silica emitters (New Objective, 75 µm inner diameter), packed in-house with reversed-phase ReproSil-Pur C18 AQ 1.9 µm resin (Dr. Maisch). A total of 0.5 µg of peptides were loaded on the column and eluted for 115 min using a segmented linear gradient of 5% to 95% solvent B. (0 min: 5% B; 0-5 min -> 5% B; 5-65 min -> 20% B; 65-90 min ->35% B; 90-100 min -> 55% B; 100-105 min ->95% B, 105-115 min - >95% B) (solvent A 0% ACN, 0.1% FA; solvent B 80% ACN, 0.1%FA) at a flow rate of 300 nL/min. Mass spectra were acquired in data-dependent acquisition mode using the TOP15 method. MS spectra were acquired in the Orbitrap analyzer with a mass range of 300–1750 m/z at a resolution of 70,000 FWHM and a target value of 3×106 ions. Precursors were selected with an isolation window of 1.3 m/z. HCD fragmentation was performed with a normalized collision energy of 25. MS/MS spectra were acquired with a target value of 105 ions at a resolution of 17,500 FWHM, a maximum injection time of 55 ms and a fixed first mass of m/z 100. Peptides with a charge of +1, greater than 6, or with an unassigned charge state, were excluded from fragmentation for MS2. Dynamic exclusion for 30s prevented repeated selection of precursors.

### MS data analysis

For MS data analysis, the raw data were processed using MaxQuant software (version 1.5.7.4, http://www.maxquant.org/) (Cox & Mann, 2008) with label-free quantification (LFQ) and iBAQ enabled (Tyanova et al., 2016). MS/MS spectra were searched by the Andromeda search engine against a combined database containing the sequences of *N. benthamiana* or *Z. mays*, along with sequences of 248 common contaminant proteins and decoy sequences. The search parameters included trypsin specificity with a maximum of two missed cleavages, a minimal peptide length of seven amino acids, and fixed modifications of carbamidomethylation of cysteine residues. Variable modifications included the oxidation of methionine and protein N-terminal acetylation. Peptide-spectrum-matches and proteins were retained if they were below a false discovery rate of 1%.

Statistical analysis of the MaxLFQ values was carried out using Perseus (version 1.5.8.5, http://www.maxquant.org/). Quantified proteins were filtered for reverse hits and hits ‘identified by site’ and MaxLFQ values were log2 transformed. Samples were grouped by condition, and only proteins with two valid values in one of the conditions (for *N. benthamiana*) or three valid values in one of the conditions (for *Z. mays*) were retained for subsequent analysis. Two-sample t-tests were performed using a permutation-based FDR of 5%. Alternatively, quantified proteins were grouped by condition, and only hits with three valid values in one of the conditions (for *N. benthamiana*) or four valid values in one of the conditions (for *Z. mays*) were retained. Missing values were imputed from a normal distribution, with a 1.8 downshift applied separately for each column. Volcano plots were generated in Perseus using an FDR of 5% and an S0 value of 1. The Perseus output was exported and further processed using Excel.

### Co-immunoprecipitation in *N. benthamiana*

*N. benthamiana* plant leaves were infiltrated with *Agrobacterium* carrying the designated constructs. At 2 days post-infiltration (dpi), the infiltrated leaves were harvested and ground into powder using liquid nitrogen. The ground powder was then mixed with protein extraction buffer (50 mM Tris-HCl pH 7.5, 150 mM NaCl, 2 mM EDTA, 10% glycerol, 1% [v/v] Triton-100, protease inhibitor cocktail (Rothe. Cat. No. 04693159001)) and incubated on ice for 30 min with inversion every 10 min. The samples were centrifuged at 4°C for 20 min at 13,000 g, and the resulting supernatant was transferred to a new tube and centrifuged again under the same conditions.

For co-immunoprecipitation assays, 1 mL of the protein supernatant was incubated with 10 μL of magnetic GFP-Trap beads (Chromotek, Cat. No. gtma) or magnetic Myc-Trap beads (Chromotek, Cat. No. yta) at 4°C with constant rotation for 1 h. The beads were collected using a magnetic stand, washed three times with extraction buffer, and then three times with a washing buffer (50 mM Tris-HCl pH 7.5, 150 mM NaCl, 2 mM EDTA, 10% glycerol, and 5 mM dithiothreitol). The beads were resuspended in 80 μL of 1x SDS-loading buffer and boiled at 95°C for 5-10 minutes. The samples were subjected to Western blotting analysis using an anti-GFP antibody (Roche, Cat. No. 11814460001) at a dilution of 1:1,000 and an anti-Myc antibody (Sigma, Cat. No. M4439) at a dilution of 1:5,000.

### RNA preparation and RNA-seq analysis

Maize seedlings were infected with *U. maydis* SG200, Δ*Tip6*, or Δ*Tip6*-*Tip6EARm*-2xHA, and the infected seedlings were harvested at 3 dpi for total RNA extraction. The infected leaves were ground into powder using liquid nitrogen and total RNA was extracted using TRIzol Reagent (Thermo Fisher Scientific, Cat. No. 15596018) following the manufacturer’s protocol. DNA removal was performed using the Turbo DNA-Free^TM^ Kit (Invitrogen, Cat. No. AM2238) according to the manufacturer’s instructions. The samples were collected with three independent replicates and subjected to library construction using Illumina NovaSeq 6000 (Illumina) at Novogene (Cambridge, UK). Trim galore (Martin, 2011) was used to remove low-quality reads and adaptors from the RNA-seq sequences. The trimmed sequences were then mapped to the *Zea mays* B73 reference genome version 5 (Schnable et al., 2009) using STAR, v. 2.7.0e (Dobin et al., 2013). Gene counts were calculated using Feature Counts (Liao et al., 2014), and genes with counts below 25 in 9 samples were removed, resulting in 29537 genes kept for analysis. Differentially expressed genes (DEGs) analysis was performed using DESeq2 (Love et al., 2014) and edgeR (Robinson et al., 2010) with cutoff criteria of absolute fold-change ≥ 1 and p-value < 0.05. The DEGs identified by both DESeq2 and edgeR were subjected to Gene Ontology (GO) term enrichment analysis using ShinyGO v.0.06 (http://bioinformatics.sdstate.edu/go65/) (Ge et al., 2020).

### Recombinant protein purification and pull-down assay

The transformed *E. coli* strains carrying the designated constructs were cultivated in dYT medium at 30°C until reaching an optical density (OD) of 0.6. Protein induction was performed by adding 0.1 mM isopropyl β-D-1-thiogalactopyranoside (IPTG) and incubating at 18°C for 16-20 h. The cell cultures were collected by centrifugation at 4000 rpm for 20 min at 4°C. The cell pellets were resuspended in lysis buffer (0.2 M NaH2PO4/0.2 M Na2HPO4 pH 6.4,150 mM NaCl, 10 mM imidazole, pH 6.4, 100 μg/ml Lysozyme, Protease inhibitor tablets), and the proteins were extracted by sonication.

The GST-Tip6ΔSP-mCherry and GST-mCherry recombinant proteins were purified using glutathione sepharose 4B beads (GE Healthcare, Cat. No. 17075601) following the product protocol. The GST tag was removed using PreScission Protease (Thermo Fisher Scientific, Cat. No. 88946). The His-ZmTPL2^N^-His recombinant protein was purified using Pierce Ni-NTA resin (QIAGEN, Cat. No. 30210) according to the product protocol. All purified recombinant proteins were further subjected to size exclusion chromatography (SEC) column and the respective protein fractions were collected. The eluted proteins were analyzed using Coomassie Brilliant Blue (CBB) stained SDS PAGE gels and western blots.

For the co-immunoprecipitation assay, the purified recombinant mCherry or Tip6ΔSP-mCherry proteins were separately mixed with purified His-ZmTPL2^N^-His proteins and incubated with pre-washed magnetic mCherry-Trap beads (Chromotek, Cat. No. rtmak) in wash buffer (0.2 M NaH2PO4/0.2 M Na2HPO4, pH 6.4,150 mM NaCl, Protease inhibitor tablets) at 4°C for 1 hour with rotation. The beads were collected using a magnetic stand and washed 5-7 times with wash buffer. Next, 100 μL of 1x SDS loading buffer was added, and the samples were boiled for 10 min. The supernatant from the boiled protein samples was analyzed using SDS-PAGE and Western blotting. Immunoblotting was performed using anti-His antibody (Thermo Fisher, Cat. No. MA1-21315) at a dilution of 1:10,000, anti-mCherry antibody (Cell Signaling Technology, Cat. No. 6g6) at a dilution of 1:3,000, and anti-mouse IgG HRP as the second antibody (Cell Signaling Technology, Cat. No. 7076S) at a dilution of 1:3,000.

### Data availability

The supporting data for the findings of this study, not included directly in the paper or its supplementary information files, can be obtained from the corresponding authors (GD) upon reasonable request. The RNAseq raw data are publicly accessible in the NCBI Gene Expression Omnibus, under accession number GSE234067. Additionally, the mass spectrometry proteomics data have been deposited in the ProteomeXchange Consortium via the PRIDE (Perez-Riverol et al., 2022) partner repository with the dataset identifier PXD042605.

### Accession numbers

In this study, Gene sequences were obtained from maizeGDB (https://www.maizegdb.org//) database. The accession numbers of genes are as follows: ZmTPL1 (Zm00001eb127680), ZmTPL2 (Zm00001eb011010), ZmTPL3 (Zm00001eb415530), ZmTPL4 (Zm00001eb398420).

## Acknowledgments

We express our gratitude to Anne Harzen for her valuable assistance in MS sample processing, and to Shan Gao for his expertise in mapping the raw RNA-seq data to count files. We acknowledge the funding received from the European Research Council (ERC) under the European Union’s Horizon 2020 research and innovation programme (grant agreement No 771035), as well as the support from the Deutsche Forschungsgemeinschaft (DFG, German Research Foundation) under Germany’s Excellence Strategy-EXC-2048/1-Project ID: 390686111. LH acknowledges financial support provided by the China Scholarship Council (Grant No. 201806320134).

## Author contributions

GD, LH and BÖ designed the research. LH, BÖ, MK, MK, DH, and AD conducted the experiments. LH performed the RNA-seq data differential gene analysis. SCS and HN carried out the MS and MS data analysis. LH and GD wrote the paper with contributions from the other authors.

## Supporting Information

**Fig. S1.** Co-immunoprecipitation to identify host interaction targets.

**Fig. S2.** Purification of mCherry, Tip6-mCherry, and 6xHis-ZmTPL2^N^-6xHis by size exclusion chromatography.

**Fig. S3.** Subcellular localization of ZmTPL2 in *N. benthamiana* and *Z. mays*.

**Fig. S4.** Absence of interaction between Tip6ΔEARs and ZmTPL2.

**Fig. S5.** Influence of Tip6 on the nuclear distribution pattern of ZmTPL2 in maize.

**Fig. S6.** Overview of differentially expressed gene analysis.

**Fig. S7.** Enriched transcription factors regulate DEGs.

**Table S1.** Mass spectrometry data of Tip6 in *N. benthamiana*.

**Table S2.** Mass spectrometry data of Tip6 in *Z. mays*.

**Table S3.** Comprehensive list of Tip6-regulated genes.

