## Supplemental Figures for "The fungal pathogen *Ustilago maydis* targets the maize corepressor TPL2 to modulate host transcription for tumorigenesis"

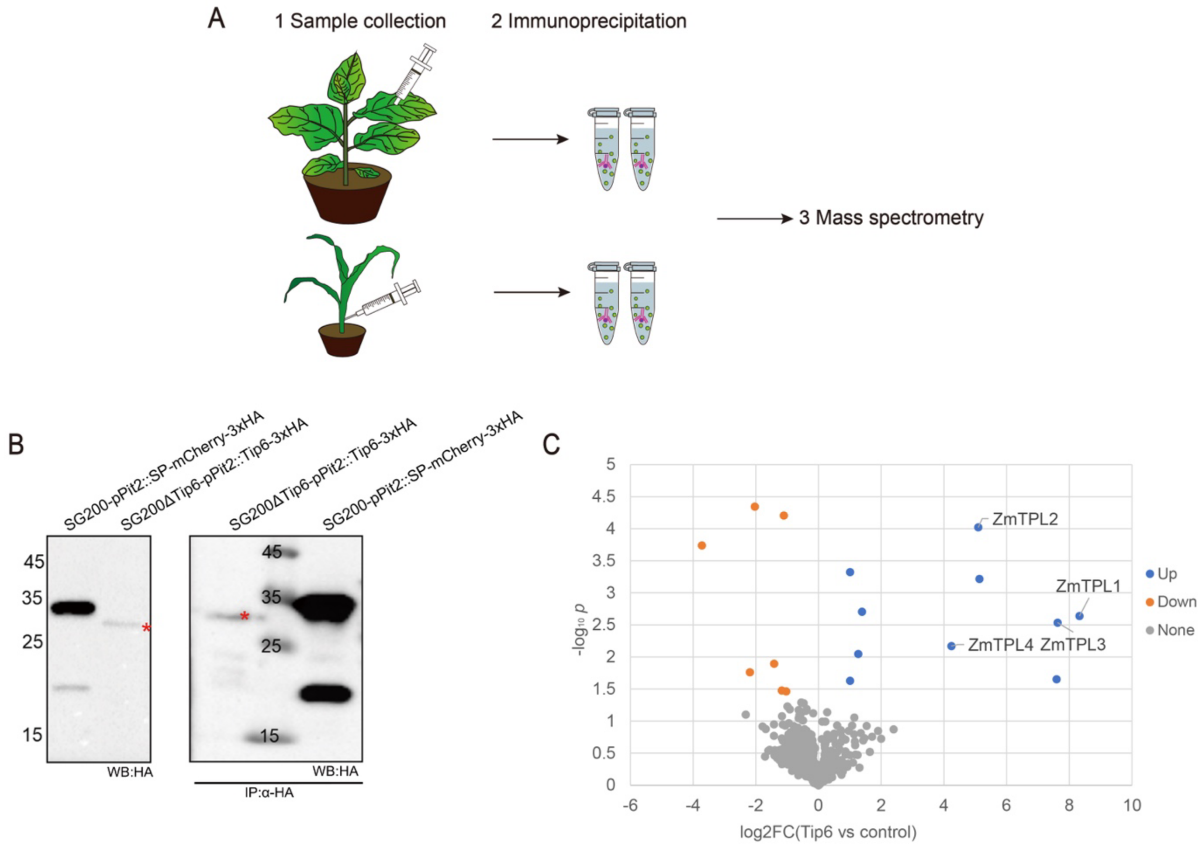


**Fig. S1. Co-immunoprecipitation to identify host interaction targets**. **A.** Workflow for the identification of host targets by mass spectrometry. (1) *N. benthamiana* was infiltrated with *Agrobacterium* strains carrying p*2x35S:: GFP* or p*2x35S::Tip6*$^{22\text{-}226}$*-GFP*. Maize leaves were infected with *U. maydis* strains expressing SG200-p*Pit2::SP-mCherry-3xHA* or SG200-p*Pit2:: Tip6-2xHA*. (2) Total protein was extracted, and a pull-down was performed according to a defined protocol (see details in the methods). (3) The pulled-down proteins were subjected to mass spectrometry. **B**. Western blot analysis of SG200-p*Pit2::SP-mCherry-3xHA* or SG200-p*Pit2:: Tip6-3xHA* in maize cell lysates and co-IP beads samples. Immunoblotting was performed using the HA antibody. The asterisk denotes the specific expected bands. **C**. Volcano plots demonstrating significant differences in the mass spectrometry of maize samples. The graph was generated in Excel. The log2 fold change (FC) represents the difference in the average label-free quantitation (LFQ) intensity of identified protein peptides in Tip6 compared to control. The *y*-axis value represents the $-\log_{10}P$-value. Each dot represents a detected protein. A log2FC >1 or -1 with a p-value < 0.05 (Student's t-test) was considered significant. Gray dots represent non-significant proteins, red dots represent down-regulated proteins, and blue dots represent up-regulated proteins. Maize TPL family genes were labeled. Control represents SG200-p*Pit2::SP-mCherry-3xHA* samples, and Tip6 represents SG200-p*Pit2::Tip6-3xHA* samples.

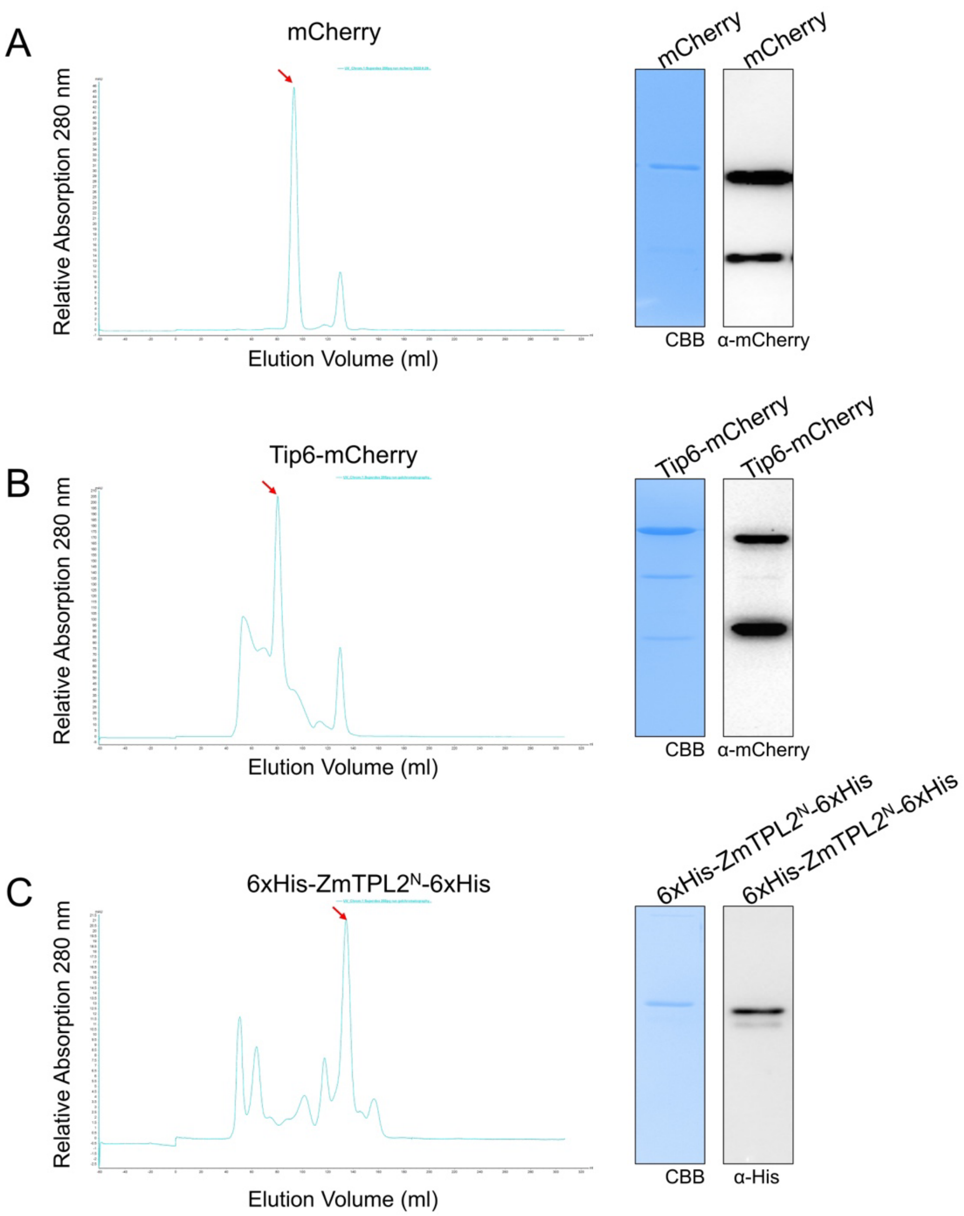


**Fig. S2. Purification of mCherry, Tip6-mCherry, and 6xHis-ZmTPL2$^{N}$-6xHis by size exclusion chromatography. A.** Purification of recombinant mCherry protein. The elution profile from the size exclusion chromatography (SEC) run with recombinant mCherry is shown on the left, with the peak corresponding to the expected size marked by a red asterisk. The right panel shows the SDS-PAGE analysis of the peak elution, which was stained with Coomassie staining and detected using mCherry antibodies. **B.** Purification of recombinant Tip6-mCherry protein. The elution profile from the SEC run with recombinant Tip6-mCherry is shown on the left, with the peak corresponding to the expected size marked by a red asterisk. The right panel shows the SDS-PAGE analysis of the peak fraction, which was stained with Coomassie and detected using mCherry antibodies. **C.** Purification of recombinant 6xHis-ZmTPL2$^{N}$-6xHis protein. The elution profile from the SEC run with recombinant 6xHis-ZmTPL2$^{N}$-6xHis is shown on the left, with the peak corresponding to the expected size marked by a red asterisk. The right panel shows the SDS-PAGE analysis of the peak fraction, which was stained with Coomassie and detected using His6 antibodies.

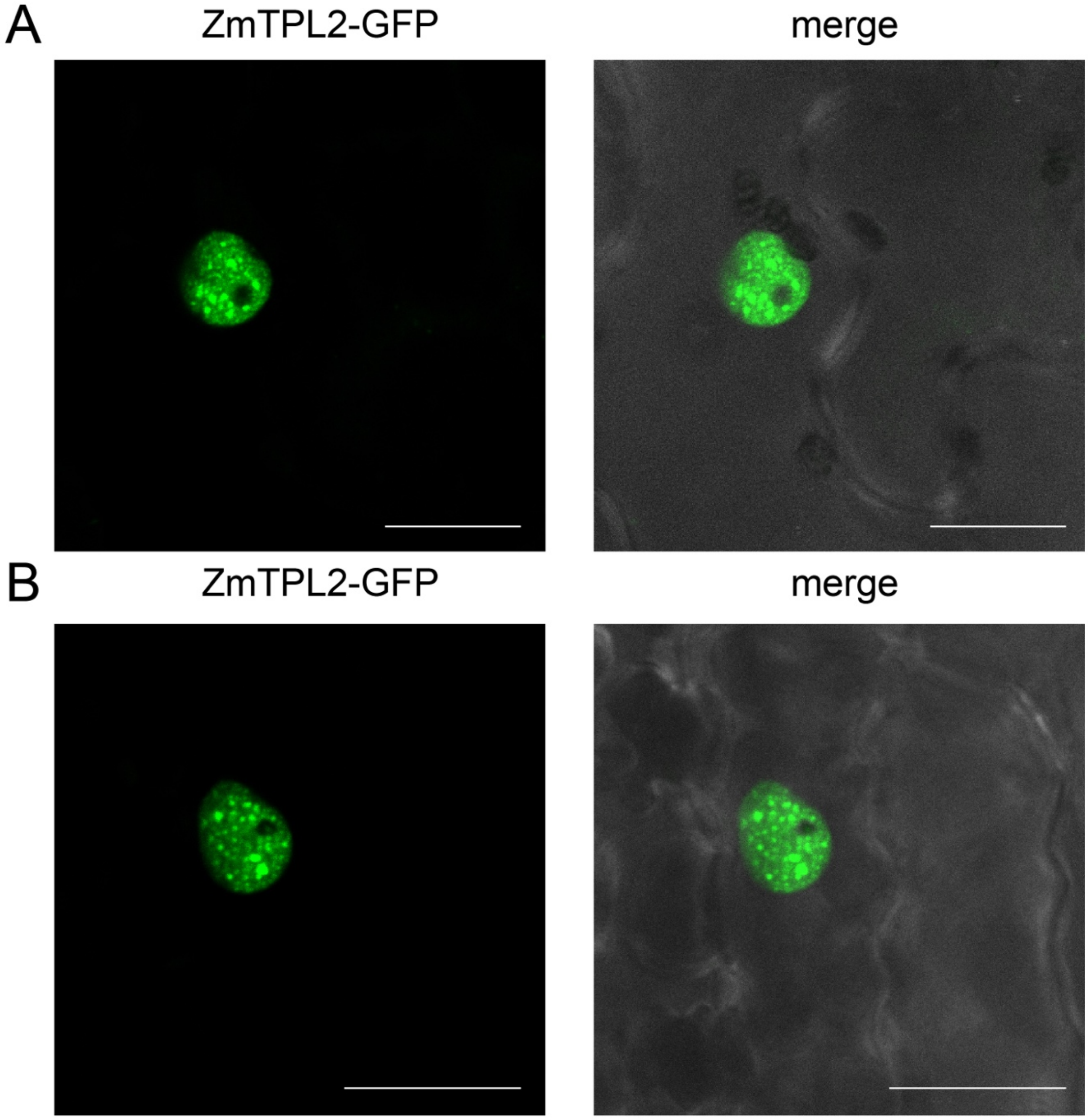


**Fig. S3. Subcellular localization of ZmTPL2 in *N. benthamiana* and *Z. mays*. A.** Confocal microscopy images showing the subcellular localization of ZmTPL2 in *N. benthamiana*. *N. benthamiana* plants were transformed with p*2x35S::ZmTPL2-GFP* using *Agrobacterium*-mediated transformation. The subcellular localization of ZmTPL2-GFP was visualized by confocal microscopy. Scale bar, 20 μm. **B.** Subcellular localization of ZmTPL2 in maize. Maize epidermal cells were transformed with p*2x35S:: ZmTPL2-GFP* using biolistic bombardment. The subcellular localization of ZmTPL2-GFP was observed 16-24 hours after transformation. Scale bar, 20 μm.

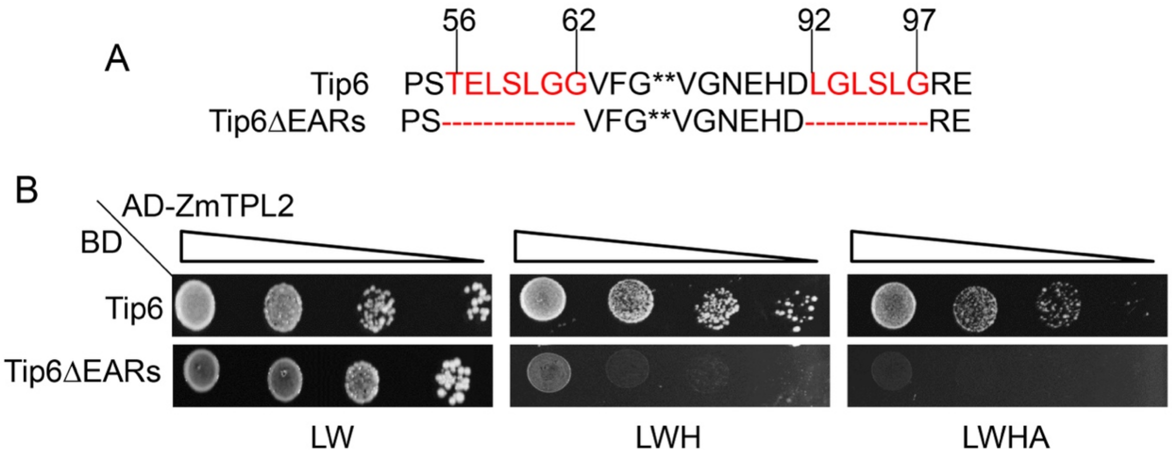


**Fig. S4. Absence of interaction between Tip6ΔEARs and ZmTPL2. A.** Schematic representation of Tip6ΔEARs protein sequence. The EAR repression domains were deleted, and the positions of the deletions are indicated by the numbers above the protein sequence. **B.** Y2H assay showing the lack of interaction between Tip6ΔEARs and ZmTPL2. The Y2H assay was performed as described in Fig. 2A and the methods section.

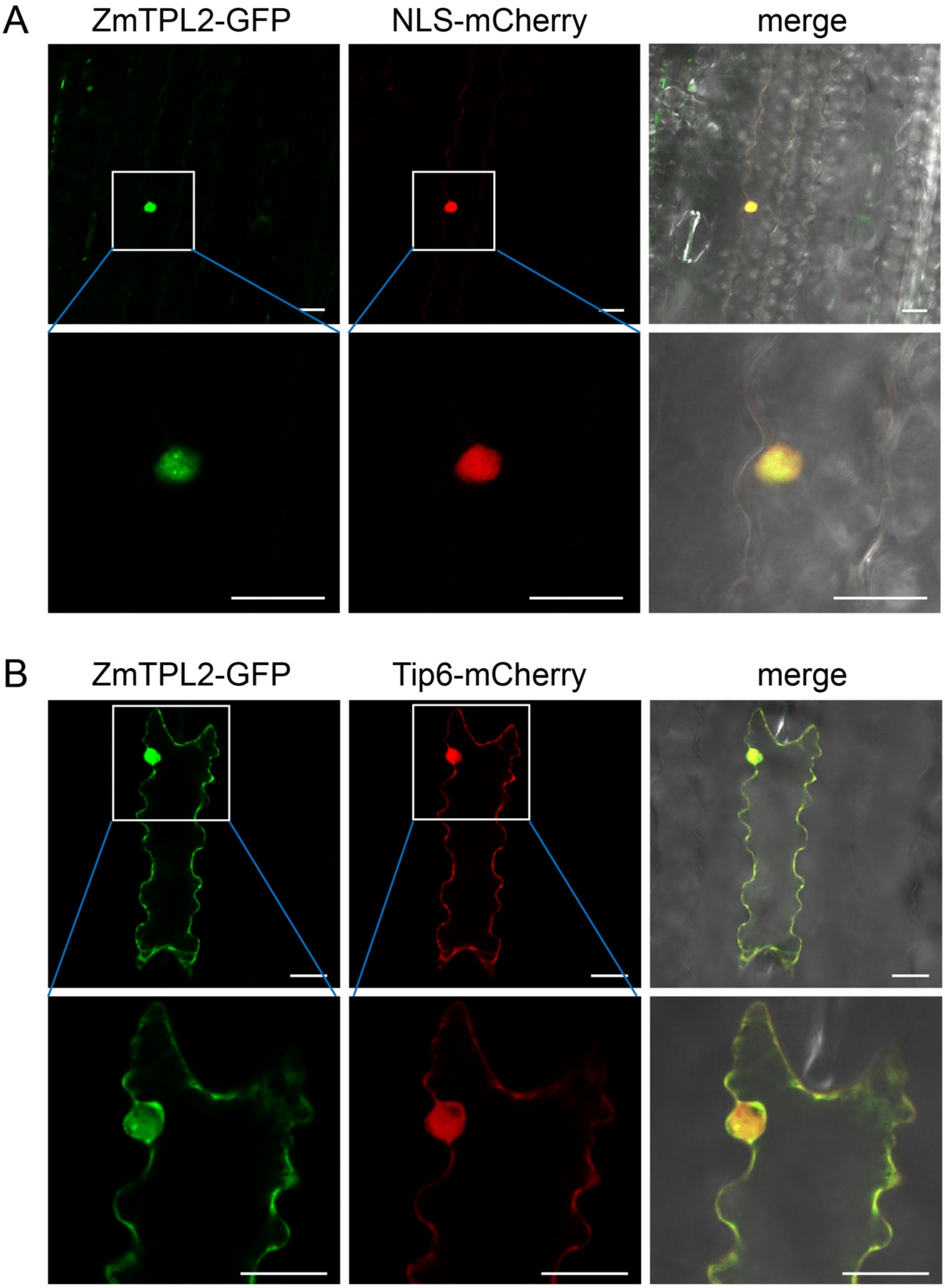


**Fig. S5. Influence of Tip6 on the nuclear distribution pattern of ZmTPL2 in maize. A.** Co-localization of ZmTPL2 with NLS-mCherry. p*2x35S::ZmTPL2-GFP* was co-expressed with p*2x35S:: NLS-mCherry* in maize epidermal cells through biolistic bombardment. The images were taken 16-24h after transformation. Scale bar, 20 μm. **B.** Co-localization of ZmTPL2 with Tip6$^{22\text{-}226}$-mCherry. p*2x35S::ZmTPL2-GFP* was co-expressed with p*2x35S::Tip6*$^{22\text{-}226}$-*mCherry* in maize epidermal cells. The images were taken 16-24h after transformation. Scale bar, 20 μm.

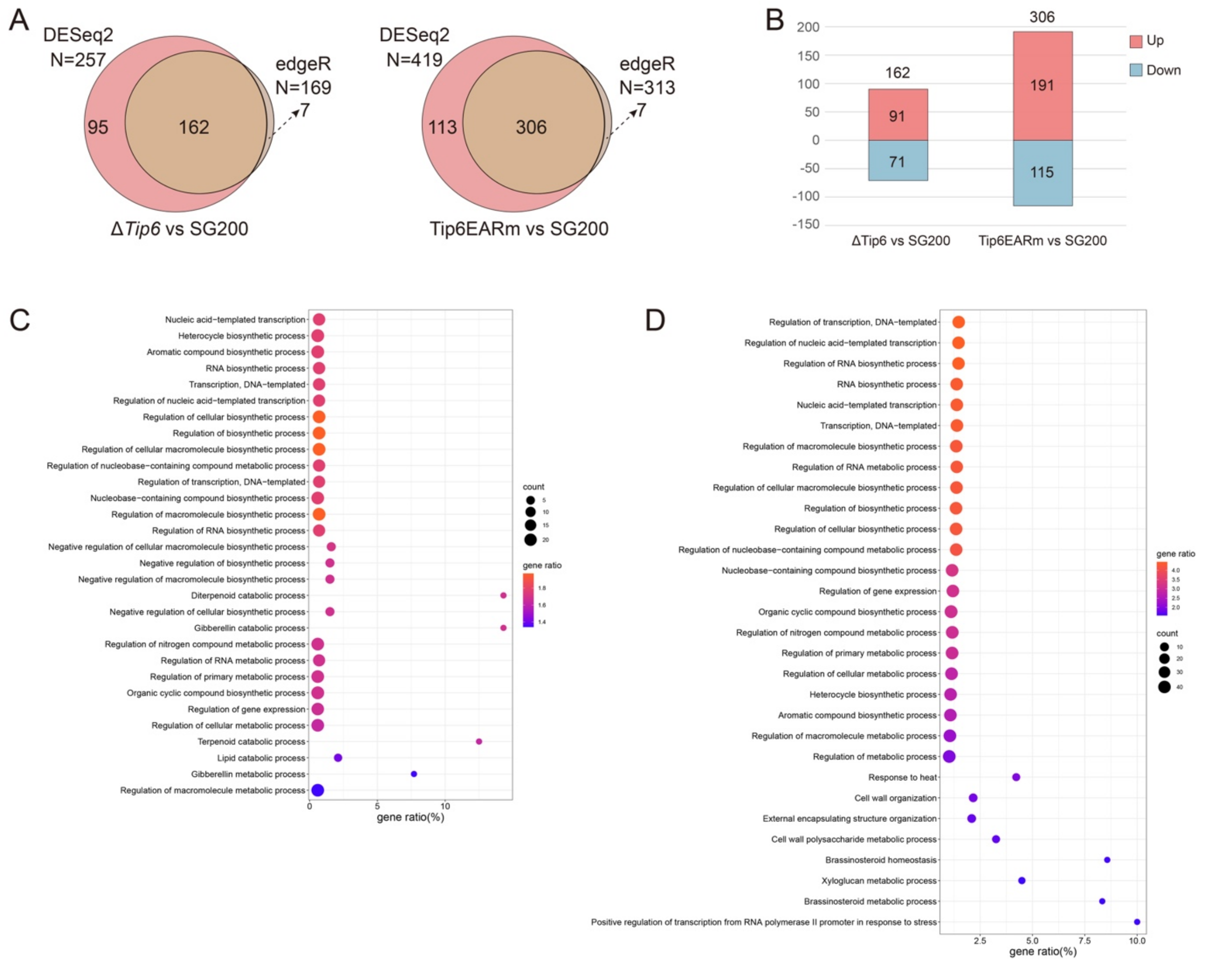


**Fig. S6. Overview of differentially expressed gene analysis. A.** Venn diagram showing the overlap of DEGs between Δ*Tip6* and SG200, and between Tip6EARm and SG200, as determined by DESeq2 and edgeR analysis. The analysis was performed using a cutoff of log2fold change > 1 and p-value < 0.05. **B.** Bar graph illustrating the number of up- and down-regulated DEGs in the Δ*Tip6* vs. SG200 and Tip6EARm vs. SG200 comparisons. **C.** GO analysis of DEGs identified in the Δ*Tip6* vs. SG200 comparison. Out of 162 DEGs, 113 were annotated with GO terms. The size of the dots represents the number of analyzed genes in the corresponding term, and the gene ratio indicates the ratio of DEGs to the total number of genes in the individual term. The gene ratio is color-coded from red to blue. **D.** GO analysis of DEGs identified in the Tip6EARm and SG200 comparison. Out of 306 DEGs, 223 were annotated with GO terms. The size of the dots represents the number of analyzed genes in the corresponding term, and the gene ratio indicates the ratio of DEGs to the total number of genes in the individual term. The gene ratio is color-coded from red to blue.

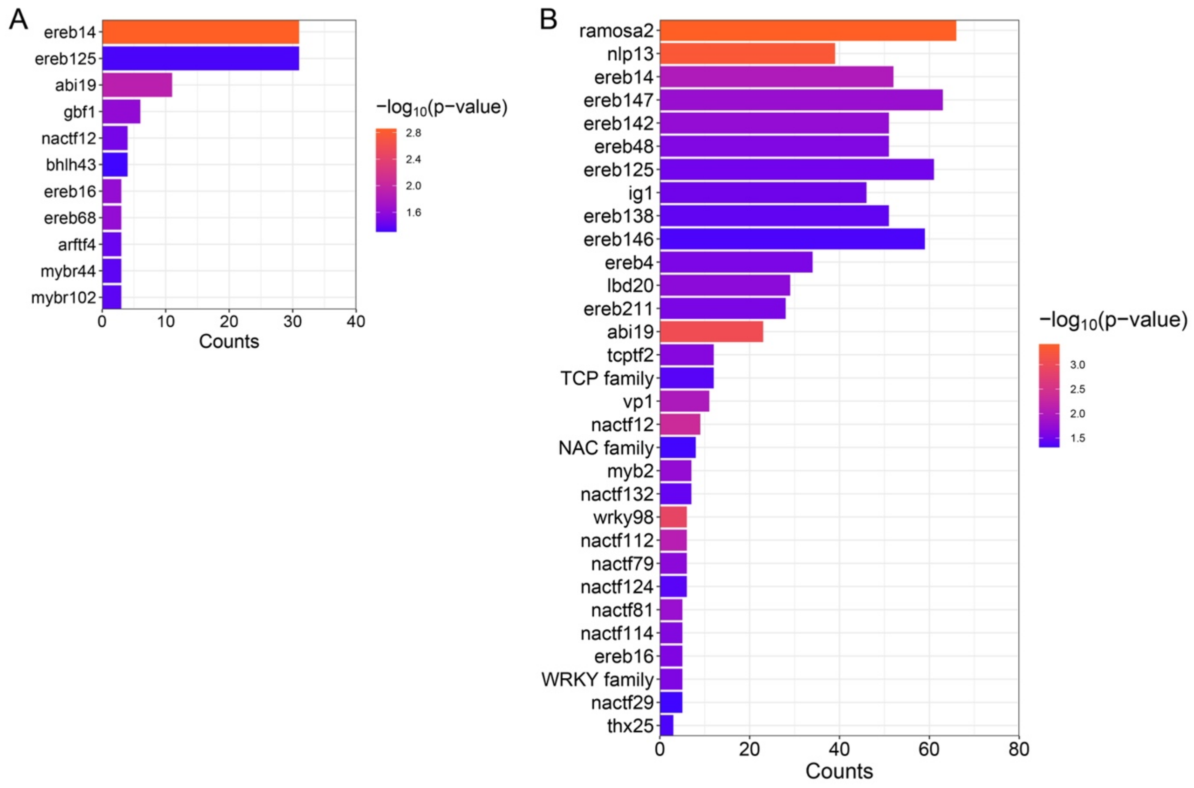


**Fig. S7. Enriched transcription factors regulate DEGs. A.** Bar graph illustrating the potential transcription factors regulating DEGs in the comparison of Δ*Tip6* and SG200. The count indicates the number of DEGs with putative binding sites for the corresponding transcription factor. The -log10 (P-value) is color-coded from blue to red, indicating the significance level of enrichment. **B.** Bar graph depicting the potential transcription factors regulating DEGs in the comparison of Tip6EARm and SG200. The count represents the number of DEGs containing putative binding sites for the corresponding transcription factor. The -log10 (p-value) is color-coded from blue to red, indicating the significance level of enrichment.
